# DESPOT: Direction-Enhanced Scoring POTentials for Scoring of Protein-Ligand Interactions

**DOI:** 10.64898/2026.03.31.714140

**Authors:** Robin Poelmans, Wout Van Eynde, Ahmed Shemy, Bence Bruncsics, Adam Arany, Yves Moreau, Arnout Voet

## Abstract

Knowledge-based potentials (KBPs) remain among the most reliable and interpretable scoring functions for protein–ligand interactions, yet most share two structural limitations. They assume that the space around each protein atom is isotropic, and their interaction-conditioned reference state cannot represent regions of space that are preferentially left empty. We introduce DESPOT (Direction-Enhanced Scoring POTentials), an all-atom anisotropic KBP that overcomes both. DESPOT classifies atoms into isotropic, axially symmetric, and fully anisotropic symmetry classes from their hybridization and bonding environment, and discretizes the surrounding interaction space using the according symmetry. By adopting a positionally averaged reference state and using a void ligand atom type, it learns, for every point around a protein atom, the probability that the point is occupied by a given ligand atom type or preferentially left empty — a ligand-independent description that naturally encodes steric exclusion. This occupancy-conditioned potential captures the precise, atom-level placement of ligand atoms; we pair it with a complementary geometry-conditioned, residue-level formulation (DESPOT-screen, in the spirit of KORP-PL) and combine the two inverse-Boltzmann scores into a consensus score, DESPOT-combo. Derived from 110,943 curated, energy-minimized complexes drawn from the CROWN database and evaluated on the CASF-2016 benchmark, DESPOT achieves competitive scoring power (Pearson *r* = 0.61), while DESPOT-combo attains best-in-class docking power (89.5% top-1 success); all anisotropic DESPOT variants significantly outperform isotropic KBPs and established empirical scoring functions in virtual screening. Anisotropy is decisive for rejecting geometrically implausible poses, and uniting the atom-level precision of DESPOT with the implicit flexibility tolerance of the residue-level score yields the most consistent performance across tasks. Because the same occupancy-conditioned potentials can be evaluated over an empty grid, DESPOT generates molecular interaction fields as well, unifying pose scoring with direction-aware binding-site characterization within a single interpretable model.

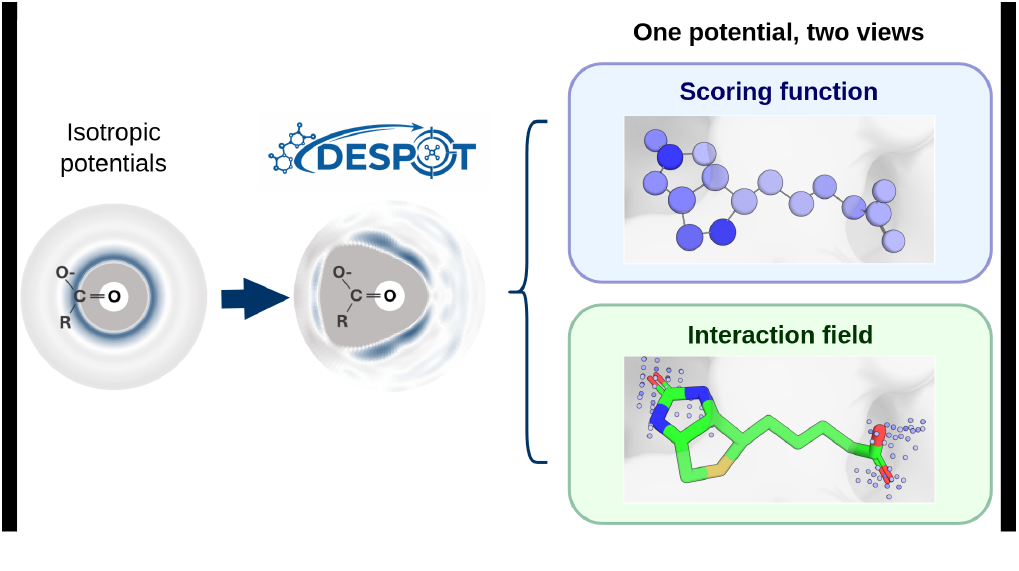

## Introduction

Predicting how a small molecule engages its protein target is central to structure-based drug discovery, and at its core lies the scoring of protein–ligand interactions: estimating how energetically favorable it is to place a given chemical group at a given position relative to a protein. Scoring functions are asked to do two related things: to judge whether a binding mode is geometrically and energetically plausible, and to rank compounds by predicted affinity for virtual screening [1]. A third, conceptually adjacent task is the construction of molecular interaction fields (MIFs), which map the three-dimensional preference of a binding site for different chemical functionalities to guide fragment placement and lead optimization [2]. Currently, MIFs are generated mostly from force-field probe calculations rather than from scoring functions, yet all three tasks pose essentially the same question of binding-site space: which chemical groups are favored where? A sufficiently expressive scoring framework could in principle serve all three — provided it can describe not only where interactions are favorable, but also where space is preferentially left empty.

Interaction scoring functions fall into three broad families. Physics-based methods derive energies from molecular-mechanics force fields, decomposing binding into electrostatic, van der Waals, and hydrogen-bonding terms [3, 4]. Empirical and machine-learning scoring functions are instead fit directly to structural and affinity data, with deep-learning models reaching strong benchmark performance [5]. Both families have well-documented weaknesses: simplified force-field functional forms struggle to capture the subtleties of shape and chemical complementarity [6], while data-driven models often generalize poorly beyond their training domain and resist interpretation [7].

Knowledge-based potentials (KBPs) sit between these extremes, pairing model flexibility with interpretability. Rather than committing to an explicit functional form, KBPs treat all intermolecular contacts within a single statistical framework and infer preferred interaction geometries directly from structural data, implicitly absorbing physicochemical effects that are difficult to parametrize by hand. Methods in this tradition, such as DrugScore [8–10], PMF [11], ASP [12] or KORP-PL [13], have remained competitive in pose scoring and virtual screening for two decades. Despite their apparent diversity, we here argue that all knowledge-based potentials for protein-ligand interactions operate on the same joint distribution *P* (*p, l*, **x**), where *p* is the protein atom type, *l* the ligand atom type, and **x** the position of *l* in the local reference frame of *p*. In particular, the above-mentioned models all compute the *interaction-conditioned* ratio *P* (**x**| *p, l*)*/P* (**x**), which measures how much more likely a position **x** is to be occupied once a *p*–*l* contact is assumed to exist [4, 9].

This formulation has two consequences that limit its reach. First, comparing positions across atom types requires consistently defined local frames, and in practice most KBPs sidestep this by assuming the space around a protein atom is isotropic. That assumption is contradicted by extensive statistical analysis of protein–ligand complexes: hydrogen bonds show pronounced angular preferences that depend on functional-group identity [14], while comparable anisotropy is reported for *π*-stacking and cation–*π* contacts [15, 16] as well as for halogen and chalcogen bonds [17]. These studies show systematic deviations from idealized valence shell electron pair repulsion (VSEPR) geometries, which are difficult to capture with physics-based functions. Second, because the ratio is conditioned on a *p*–*l* contact being present, it cannot represent the *absence* of a ligand atom: it has no means to express that a region of space is sterically forbidden. It is therefore intrinsically incompatible with MIF generation, which requires reasoning about empty space.

Two prior advances each addressed one of these limitations. KORP-PL abandons the isotropy assumption, scoring interactions in orientation-dependent frames built from protein C*α* atoms and distinguished by residue type [13]. This buys directional sensitivity and strong virtual-screening performance, but at a price: the C*α*-only representation is coarse-grained and cannot attribute interactions to specific functional groups or cofactors, and — being interaction-conditioned like the rest of the family — it still presupposes that a ligand atom occupies each scored position. It thus inherits the second limitation and can neither account for steric exclusion nor generate MIFs. Separately, DFIRE [18] replaced the interaction-conditioned ratio with a so-called “finite ideal-gas” reference state, effectively computing *P* (*l* | *p*, **x**)*/P* (*l* | *p, r*_max_): how much more likely ligand atom *l* is to be found at position **x** than in its non-interacting “ideal gas’” state far from the protein surface. Crucially, this occupancy-conditioned view is defined for *every* point in space, whether or not an interaction occurs there, so depletion — and hence steric exclusion — emerges naturally. The formulation was originally developed as an isotropic potential for protein structure modeling.

DESPOT (Direction-Enhanced Scoring POTentials) unifies these two ideas into what is, to our knowledge, the first all-atom anisotropic KBP for protein–ligand interactions. The combination of atom-level resolution and an occupancy-averaged reference state yields three advantages over KORP-PL: it can also capture contributions from both organic and inorganic cofactors instead of only working with standard residues; it can account for steric exclusion, making it directly applicable to MIF generation; and finally, the smaller atom-level reference frames offer much higher interpretability on how interactions work. In effect, DESPOT learns, for every region of space around a protein atom, the probability that it is occupied by a particular ligand atom type or preferentially left empty — a ligand-independent, probabilistic picture of binding-site space that unifies pose scoring with directional, steric-aware site characterization. This occupancy-conditioned, atom-level score is the core of DESPOT, but it is not the only conditioning of *P* (*p, l, x*) worth exploiting. Because the same joint distribution also admits a geometry-conditioned, residue-level reference state — the formulation underlying KORP-PL — we derive it here as a complementary potential, DESPOT-screen, using the same training data, atom typing, and discretization for a controlled comparison. The two scores capture different signals: the atom-level DESPOT resolves the precise placement of individual ligand atoms, whereas the coarser residue-level DESPOT-screen is largely blind to exact side-chain rotamers and thereby tolerates the conformational adjustments a flexible receptor would make. Summing them yields a consensus score, DESPOT-combo, that combines these type- and geometry-conditioned views and, as we show below, delivers the most consistent performance across benchmark tasks.

Capturing these directional preferences requires reference frames adapted to local atomic geometry. Central to DESPOT is the observation that atoms exhibit different local symmetries depending on hybridization state and bonding connectivity. Inspired by the local reference frames of Schuetz et al. [19], we classify atoms into isotropic, axially symmetric, or fully anisotropic symmetry classes, each with its own geometric discretization (Figure 1), and describe interactions in atom-centered frames using spherical shells, sectors, or voxels as appropriate [20]. This adaptive discretization preserves the simplicity of classical KBPs while extending their geometric expressiveness to three dimensions.

**Figure 1:**
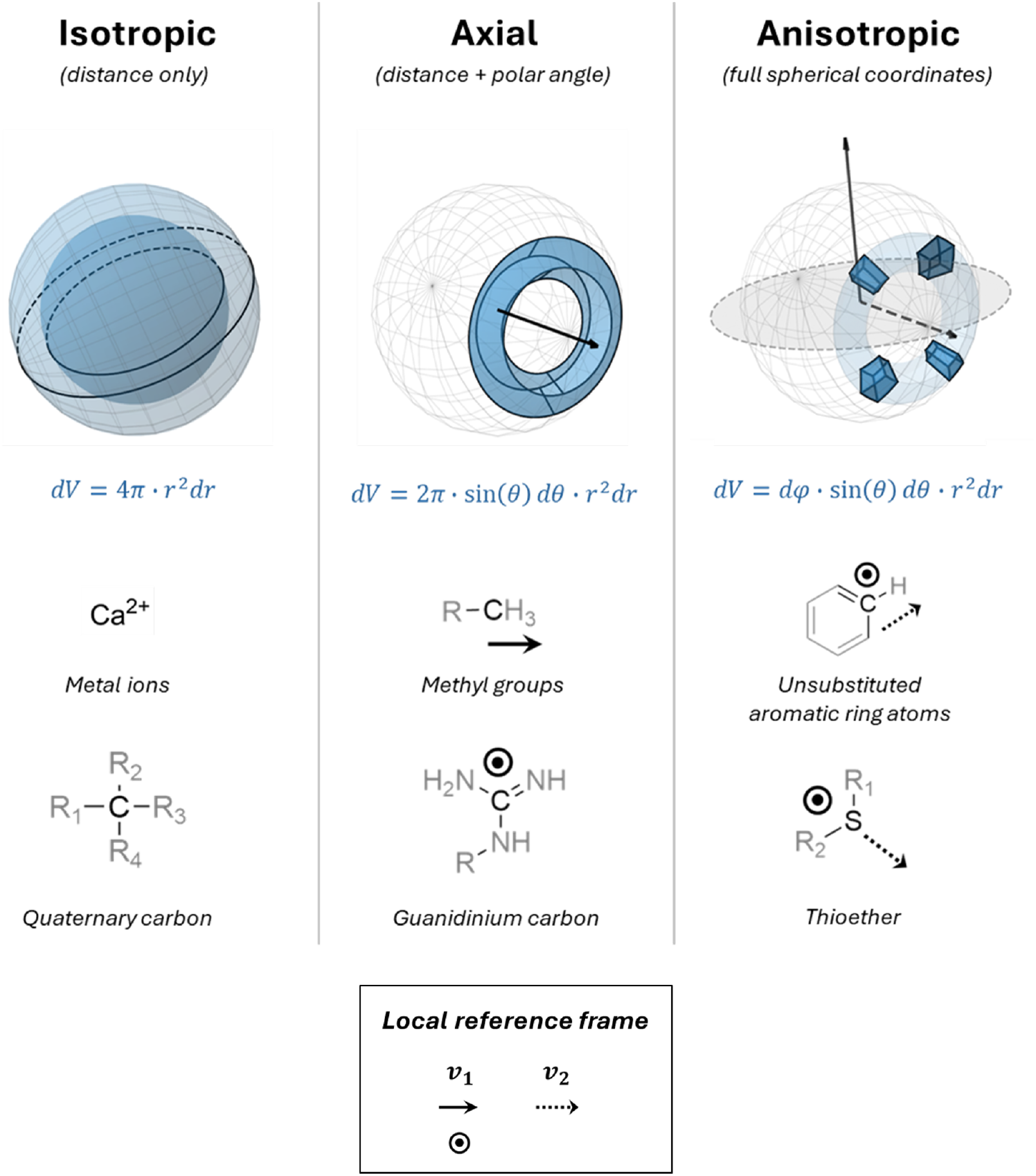
Symmetry-based geometric discretization for anisotropic interaction modeling. Atoms are classified into three symmetry classes based on hybridization state and bonding connectivity, each with a corresponding geometric discretization scheme. Approximately isotropic atoms (e.g., metal ions, quaternary carbons) exhibit spherical symmetry; interactions are binned in concentric spherical shells depending only on radial distance *r*, with differential volume *dV* = 4*π · r*^2^*dr*. Axially symmetric atoms (e.g., methyl groups, guanidinium carbon) possess rotational symmetry about a single axis; interactions are binned in spherical sectors using distance *r* and polar angle *θ*. Fully anisotropic atoms (e.g., unsubstituted aromatic ring carbons, thioether sulphur) require full three-dimensional characterization; interactions are binned in spherical voxels using *r, θ*, and azimuthal angle *ϕ*. Representative functional groups are shown for each symmetry class.

This expressiveness comes at a statistical cost. Moving from radial to angular discretization sharply increases the number of parameters to estimate, demanding training data that are simultaneously large and structurally consistent — a combination existing resources did not provide. Heavily curated collections such as HiQBind achieve high structural quality through extensive preprocessing but remain modest in size [21], whereas the large-scale PLINDER initiative offers unprecedented coverage but only minimal quality filtering [22]. To close this gap between rigor and scale, we recently created CROWN (Curated Repository Of Well-resolved Non-covalent interactions), a machine-learning-ready dataset of 178,263 high-quality protein–ligand complexes produced through systematic correction, protonation, and restrained energy minimization [23].

In this work we derive DESPOT potentials from CROWN and evaluate them on the CASF-2016 benchmark, which assesses the scoring, ranking, docking, and screening power of scoring functions [24]. We find that anisotropic potentials substantially outperform their isotropic counterparts — most clearly in penalizing geometrically implausible poses — and perform on par with established empirical scoring functions. We further analyze how restrained energy minimization and rigorous train–test splitting affect performance, and show that the learned interaction profiles recover systematic directional preferences for hydrogen bonds, aromatic interactions, and halogen bonds that go beyond idealized geometric models. Together, these results establish DESPOT as a scalable framework for direction-aware modeling of protein–ligand interactions that unifies pose scoring with a ligand-independent, probabilistic characterization of binding sites.

## Methods

### Theory: Direction-Enhanced Scoring POTentials

Most KBP algorithms operate on the same joint probability distribution (JPD) *P* (*p, l*, **x**), with protein atom type *p*, ligand atom type *l*, and **x** the position of *l* in the local reference frame of *p*. This JPD is discrete along *p* and *l* but continuous across its spatial axes.

The conceptual move at the heart of DESPOT is to invert the usual conditioning. Rather than asking *where a ligand atom lies given that a contact exists* (the interaction-conditioned ratio *P* (*x* | *p, l*)*/P* (*x*)), DESPOT asks *which ligand atom type, if any, occupies a given point in space*, through the occupancy-conditioned ratio *P* (*l* | *p, x*)*/P* (*l* | *p*). Because this quantity is defined at every point around a protein atom regardless of whether an interaction occurs there, the same model describes both favorable contacts and empty space. This change allows DESPOT to serve both pose scoring and MIF generation.

In the following we show how the JPD can be constructed from any dataset of protein–ligand complexes through a sequence of steps: atom typing of protein and ligand atoms; construction of local reference frames; binning of pairwise interaction counts; volume normalization and symmetry expansion of the spatial bins; heat-kernel smoothing; addition of a void ligand density; and conversion to conditional probabilities and pairwise pseudo-energies.

### Atom typing and local reference frame construction

Atom typing determines the resolution at which interaction preferences are learned, and therefore controls the balance between chemical specificity and statistical robustness. DESPOT closely follows the *fconv* atom typing scheme used in DrugScoreX, constructing environment-dependent atom types from SYBYL MOL2 base types refined by first- and second-degree heavy-atom bonding environments [9, 25]. First-degree descriptors encode the number and elemental identity of bonded heavy atoms, hydrogen count, and hybridization; second-degree descriptors capture the coordination state of neighboring atoms, separating chemically distinct functional groups that share identical first-shell connectivity.

The principal extension relative to *fconv* is that DESPOT co-defines atom type and symmetry class: whereas *fconv* may assign a common label to atoms differing only in heavy-atom-neighbor count (e.g., un-substituted and substituted aromatic carbons are both labeled *C*.*ar*6), DESPOT treats these as distinct types, because heavy-atom coordination determines an atom’s symmetry classification and hence the geometric discretization of its surrounding interaction space. Linking chemical environment and geometric discretization intrinsically avoids retrofitting directional information onto a typing system not designed to encode it. The resulting atom typing scheme used by DESPOT defines 67 protein atom types and 145 ligand atom types (atoms falling outside these definitions are omitted). The full set of refinement rules parallels that of *fconv* and is detailed in the Supporting Information (Table S1, Figure S2).

Each atom type is assigned to one of three symmetry classes, which govern how the surrounding interaction space is discretized (Figure S1); the complete mapping from hybridization and heavy-atom-neighbor count to symmetry class is given in Figure 2.

**Figure 2:**
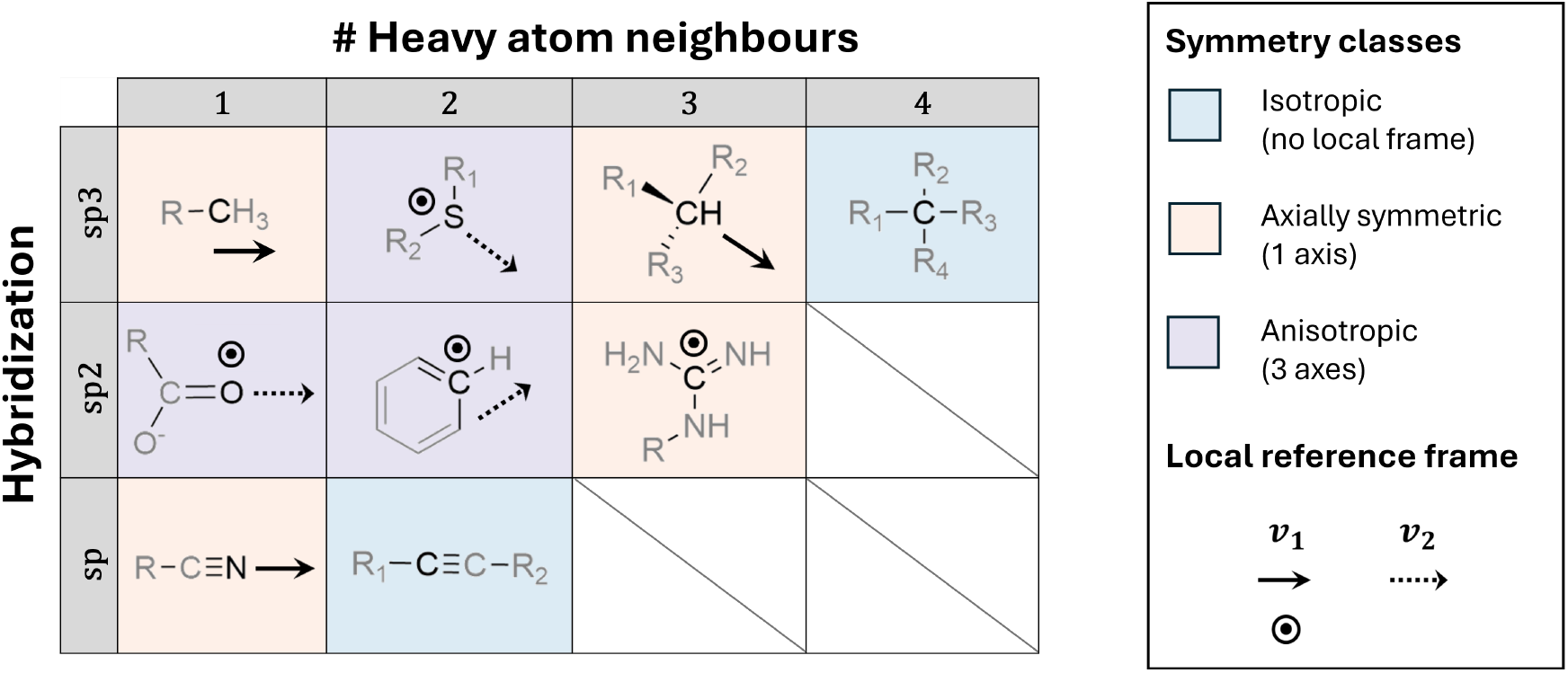
Local reference frame assignment by atom symmetry class. Classification of atoms into symmetry classes based on hybrid9.ization state and number of heavy-atom neighbors. Each combination is assigned to the isotropic (blue), axially symmetric (orange), or fully anisotropic (purple) symmetry class, which determines the geometric discretization used for binning interactions. For axially symmetric and anisotropic atoms, local reference frame axes (**v**_**1**_, **v**_**2**_) are constructed to align with chemically meaningful directions: bond axes for axially symmetric atoms, plane normals and in-plane bond vectors for anisotropic atoms. Representative molecular fragments illustrate each category.

- **Isotropic atoms** (e.g., ions, or sp^3^ atoms with four heavy-atom neighbors) are treated as spherically symmetric, with interactions binned into radial shells only.
- **Axially symmetric atoms** (e.g., sp- or sp^3^-hybridized atoms with a single heavy-atom neighbor) are assigned a single signed reference axis **v**_1_, enabling discretization into spherical sectors that resolve direction along the bond axis while averaging over rotation about it. Here **v**_1_ is the unit vector from the bonded neighbor toward the atom of interest; for sp^3^ atoms with three heavy-atom neighbors, **v**_1_ is instead the unit vector from the centroid of the three neighbors to the central atom, approximating the remaining tetrahedral lobe.
- **Fully anisotropic atoms** (sp^2^ centers, and other planar or low-coordination atoms such as thioether sulfur) are assigned a complete orthonormal basis (**v**_1_, **v**_2_, **v**_3_), enabling full spherical discretization: **v**_1_ is the normal to the local plane defined by the atom and two of its neighbors; **v**_2_ lies along the bisector of the two in-plane bond vectors (or, for a single heavy-atom sp^2^ neighbor, along the bond axis, with the plane inferred from that neighbor’s geometry); and **v**_3_ completes the right-handed basis.

The three-tier scheme is a deliberate approximation. In principle the local environment of almost any atom becomes unique — and therefore fully anisotropic — once third-degree neighbors are considered, but such granularity would yield an unmanageable number of atom types, each too sparsely sampled to support reliable potentials. The scheme adopted here balances the dominant geometric features that shape directional preferences against statistical power per atom type.

### Coordinate transformation and interaction binning

For each protein–ligand atom pair, the displacement **d** from the protein atom to the ligand atom is expressed in spherical coordinates within the protein atom’s local frame. The radial coordinate is *r* = ∥**d**∥ in all cases.

The angular coordinates depend on symmetry class, because the reference axes differ in whether their sign is physically meaningful.

For **axially symmetric** atoms, **v**_**1**_ is a signed bond axis, so the polar angle retains its full range and no azimuthal angle is used (Eq. 1):

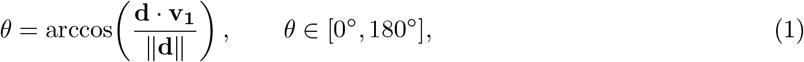

For **fully anisotropic** atoms, **v**_**1**_ is a plane normal with no intrinsic sign; the right-handed construction then ties the sign of **v**_**3**_ to that of **v**_**1**_, while the in-plane bisector **v**_**2**_ remains signed. Assuming local *C*_2*v*_ symmetry of the binding environment — the molecular plane and the plane containing **v**_**1**_ and **v**_**2**_ both act as mirrors — we fold both *θ* and the azimuthal angle *ϕ* by taking absolute values (Eq. 2):

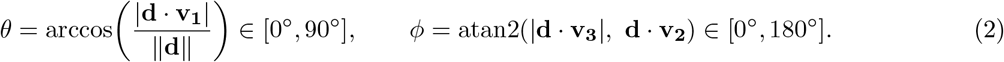

The two-argument arctangent atan2(*y, x*) returns the angle of (*x, y*) from the positive *x*-axis, preserving quadrant information; retaining the sign of **d v**_**2**_ distinguishes the bisector and anti-bisector hemispheres. This *C*_2*v*_ folding is exact for symmetric centers (e.g., aromatic CH with two equivalent ring neighbors) and approximate for asymmetric ones, where the two sides along **v**_**3**_ are not strictly equivalent.

Following Neudert and Klebe [9], interactions were recorded within a radial range of 1–6 Å (Δ*r* = 0.1 Å). Axially symmetric atoms were binned in (*r, θ*) and fully anisotropic atoms additionally in *ϕ*, at 3.0° angular resolution. The angular bins of each radial shell are laid out on a Driscoll–Healy (DH2) grid [26] an equiangular sampling of the spherical surface *S*^2^ for which the spherical-harmonic transform is exact so that each shell can be smoothed analytically in the spectral domain (below) rather than by real-space convolution.

### From interaction counts to interaction density

To enable heat-kernel smoothing and the void density calculation below, raw counts were first converted to a density by normalizing each bin for its volume *V*_*b*_, which depends on symmetry class and on the multiplicity introduced by treating reference axes as unsigned (Eq. 3):

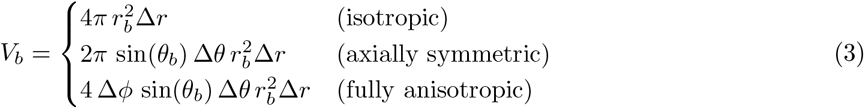

where **x**_**b**_ = (*r*_*b*_, *θ*_*b*_, *ϕ*_*b*_) is the bin center and the factor of 4 in the anisotropic case accounts for the four equivalent voxels identified by the *C*_2*v*_ folding of *θ* and *ϕ*. For uniform downstream processing, these reduced-domain densities were expanded onto a full-sphere voxel grid by copying bin densities to their symmetry-equivalent voxels.

### Smoothing of the interaction density

Volume-normalized, symmetry-expanded densities were smoothed to suppress sampling noise from finite data and discretization; without smoothing, sparsely populated bins yield unstable estimates and exaggerated energies after the inverse-Boltzmann step. Smoothing over a sphere requires separate radial and angular treatments. Radial smoothing is a Gaussian convolution along *r*, but naïve Gaussian convolution in (*θ, ϕ*) is not isotropic on *S*^2^, which lacks the translational invariance assumed by standard convolution.

Angular smoothing is therefore formulated intrinsically. Any square-integrable function on *S*^2^ admits a spherical-harmonic expansion (Eq. 4)

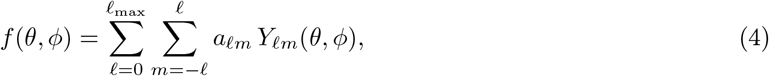

where the degree *l* indexes angular frequency: low *l* describes broad structure, high *l* fine detail. Smoothing is applied as a single multiplicative filter in the spectral domain, depending only on *l* (Eq. 5):

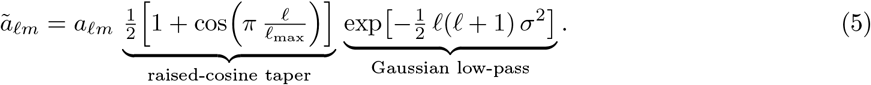

The Gaussian factor performs isotropic heat-kernel smoothing on the sphere with bandwidth *σ* (in radians), attenuating the high-frequency components that largely reflect sampling noise [27]. The raised-cosine factor is a spectral window that tapers the coefficients smoothly to zero as *l* → *l*_max_: because the expansion is truncated at a finite *l*_max_ (determined by the binning resolution), this suppresses the Gibbs (ringing) oscillations that an abrupt cutoff would otherwise introduce [28, 29]. In practice, radial smoothing was applied first with *σ*_*r*_ = 0.1 Å (following Neudert and Klebe [9]), followed by spectral filtering with *σ* = 0.1 rad using SHTools [30].

### Addition of void density

To represent regions of space that preferentially remain unoccupied, a pseudo-ligand type *l*_*∅*_ (“void”) was introduced, with per-bin density chosen so that the marginal *P* (**x** | *p*) is uniform (Eq. 6):

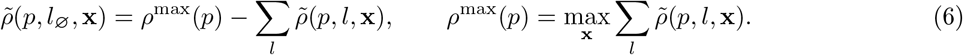

This assigns high void probability where few ligand atoms are observed, thereby encoding both steric exclusion and unfavorable interaction geometries. The void type is a reference construct and is never scored directly; it enters only through the conditional normalization below.

### From interaction density to interaction probability

After normalization of the augmented density(including *l*_*∅*_) to a JPD (Eq. 7),

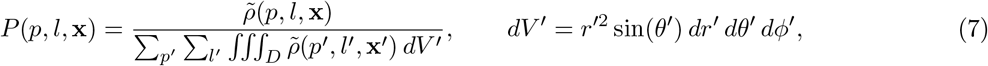

the conditional probability follows directly, with the sum over *l*^*′*^ running over all ligand types *including* the void type, so that the absence of a ligand atom is treated on the same footing as its presence (Eq. 8):

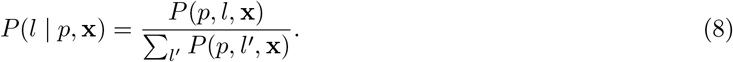

The reference distribution used in DESPOT, *P* (*l* | *p*) is the bulk composition of ligand atoms expected in the absence of interaction, and is obtained by marginalizing over spatial position. Thus *P* (*l* | *p*, **x**) is the likelihood of ligand type *l* at position **x** in the frame of protein type *p*, while *P* (*l* | *p*) is its likelihood in the non-interacting bulk.

### Pseudo-energy scoring via the inverse Boltzmann relation

Pairwise scores compare the conditional probability to the bulk reference (Eq. 9):

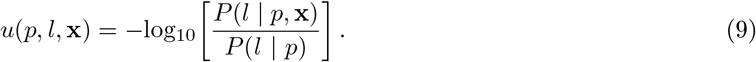

A negative *u* indicates that *l* is observed more often than expected at **x** around *p*, a favorable interaction. To guard against numerical instability in sparsely sampled bins, the probability ratio was floored at 10^*−*12^ before the logarithm, and the resulting pseudo-energies were capped to the range [−5, +5] to prevent individual bins from dominating the total score. The total score sums over all pairs within the interaction cutoff (Eq. 10),

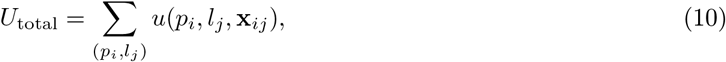

where **x**_*ij*_ is the position of ligand atom *j* in the frame of protein atom *i*.

### Training sets and ablation studies

The derivation of anisotropic interaction potentials requires substantially larger and cleaner training datasets than classical radial KBPs, because the increased dimensionality of the geometric discretization amplifies the effect of sampling noise. DESPOT was trained on a drug-like subset of the CROWN database, a curated collection of energy-minimized, high-quality protein–ligand complexes [23]. To avoid train–test leakage with CASF-2016 [24], all CROWN entries containing both a protein chain with *>* 70% sequence identity to any protein chain in CASF-2016 and *>* 70% ECFP4 Tanimoto similarity to its associated ligand were excluded. Since KBPs are trained on the interaction level, splitting on protein or ligand similarity alone is insufficient. Additional drug-likeness filters were applied: complexes containing oligopeptide or oligosaccharide ligands were discarded, and ligands were required to satisfy a quantitative estimate of drug-likeness (QED) score greater than 0.1 [31]. To reduce bias from over-represented ligands, the dataset was further restricted to a maximum of 500 complexes per CCD code, with entries selected at random above this threshold. This cap prevents frequently co-crystallized ligands from dominating the derived potentials and promotes a more balanced representation of interaction space. The final training set comprises 110,943 protein–ligand complexes.

To benchmark DESPOT against established knowledge-based potentials on equal footing, we re-implemented two reference formulations in Python: DESPOT-iso, a radial (distance-only) potential analogous to DrugScore [10], and DESPOT-screen, an anisotropic potential analogous to KORP-PL [13]. All variants were trained using the same atom-typing scheme, training data, geometric discretization, and smoothing as DESPOT. This ensures a consistent and fair comparison that isolates the effect of the probabilistic formulation from differences in training data or atom-type definitions.

The variants differ in the conditional probability they evaluate and, as a consequence, in the reference frame required to express the protein–ligand geometry (Eq. 11). DESPOT models the ligand-type distribution at a given position, *P* (*l* | *p*, **x**), relative to the spatially averaged reference *P* (*l* | *p*). DESPOT-screen, by contrast, models the spatial distribution of interactions, *P* (**x**, relative to the type-averaged *P* (**x**). Because this reference is itself a distribution over positions, it must be accumulated in a single, consistent anisotropic frame. Following KORP-PL, this frame is defined at the residue level, making the anisotropic reference *P* (**x**) well-defined. Finally, DESPOT-combo is the sum of the DESPOT and DESPOT-screen scores, combining the complementary type- and geometry-conditioned signals.

Here *p* and *l* denote protein and ligand atom types, *r* the interatomic distance, and *x* the ligand position expressed in the appropriate protein reference frame (atom-centered for DESPOT, residue-centered for DESPOT-screen).

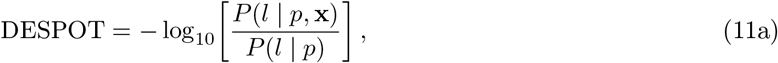

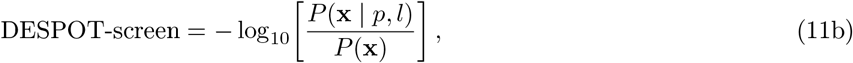

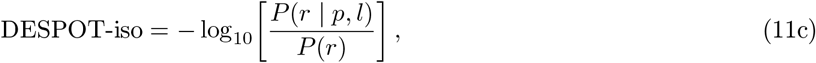

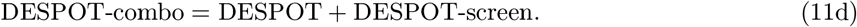

### CASF-2016 benchmark

DESPOT and all its variants were evaluated on the CASF-2016 benchmark [24]. CASF-2016 comprises 285 high-quality protein–ligand crystal complexes grouped into 57 protein clusters. The benchmark is designed to decouple pose scoring from pose generation by providing standardized ligand poses and pre-generated decoy sets, ensuring reproducibility and comparability across methods. As noted above, all PDB entries occurring in CASF-2016 were excluded from the CROWN training set during potential derivation.

For comparison, we benchmarked against a set of established scoring functions, including DrugScoreX [9], DrugScoreCSD [8], DrugScore2018 [10], ASP [12], PMF04 [11], KORP-PL [13], GlideScore-SP [32], GoldScore [33], AutoDock Vina [34], ΔVinaRF20 [35], ChemPLP [36], ChemScore [37], MOE’s GBVI-WSA-dG and GNINA (v1.3) [39]. Performance data for all methods were taken from the original CASF-2016 publication, with the exceptions of DrugScoreX and GNINA, for which the results were computed independently. For GNINA, we include three separate scores: predicted affinity (GNINA-affinity), predicted pose score (GNINA-score) and virtual screening score GNINA-VS, the product of GNINA-affinity and GNINA-score. Performance was assessed across four complementary criteria:

- **Scoring power** quantifies the ability to reproduce experimental binding affinities and is defined as the Pearson correlation coefficient between predicted scores and measured binding constants (in log units) across the 285 complexes of the benchmark.
- **Ranking power** assesses whether ligands of a common target are ordered correctly by relative affinity. For each of the 57 targets, represented by five ligands of known affinity, the Spearman rank correlation between predicted score and experimental affinity is computed and then averaged over all targets.
- **Docking power** evaluates pose selection, namely the ability to recover a near-native binding pose (within 2.0 Å RMSD of the crystallographic ligand) from a set of pre-generated decoy poses. The breadth of the binding funnel is additionally characterized by the Spearman correlation between score and RMSD.
- **Screening power** measures the discrimination of true binders from decoys in a virtual-screening setting and is reported as enrichment factors over the top 1%, 2%, and 5% of ranked compounds. CASF-2016 probes this in two directions: forward screening tests whether a target’s true ligands can be separated from decoys, whereas reverse screening tests the converse, assigning each ligand to its cognate receptor from a panel of candidate targets.

Statistical uncertainty in all performance metrics was quantified using bias-corrected and accelerated (BCa) bootstrapping with 10,000 resampling iterations [24, 40]. Performance metrics were recomputed for each bootstrap replica, yielding empirical distributions and associated confidence intervals. Pairwise comparisons between scoring functions were conducted using the non-parametric Friedman-Nemenyi test, with scoring functions considered statistically indistinguishable at *p >* 0.10, following the protocol of Su et al. [24]. For further detail on the performance tests and the reference scoring functions, we refer to Su et al. [24].

## Results and Discussion

Anisotropy matters most where binding geometry is in question. For a correctly placed ligand the directional constraints are already satisfied, so angular information adds little to its predicted affinity; for an implausible pose, that same information exposes the violated interaction geometries a distance-only model cannot see. This asymmetry (minor gains in scoring, substantial ones in pose discrimination and screening) organizes the results below. We first establish that DESPOT recovers physically meaningful directional patterns and supports both pose scoring and MIF generation, then quantify its performance on CASF-2016.

### Anisotropic interaction patterns

A major advantage of knowledge-based potentials over machine-learning scoring functions is their interpretability. Because the energy terms are fully additive, each interaction type can be examined separately, providing direct insight into the physicochemical forces governing molecular recognition. Figure 3 presents representative anisotropic interaction profiles from DESPOT, illustrating how directional preferences emerge naturally from the statistical analysis of protein–ligand complexes. (A Jupyter notebook for the visualization of pairwise interaction potentials is provided in the accompanying data repository.)

**Figure 3:**
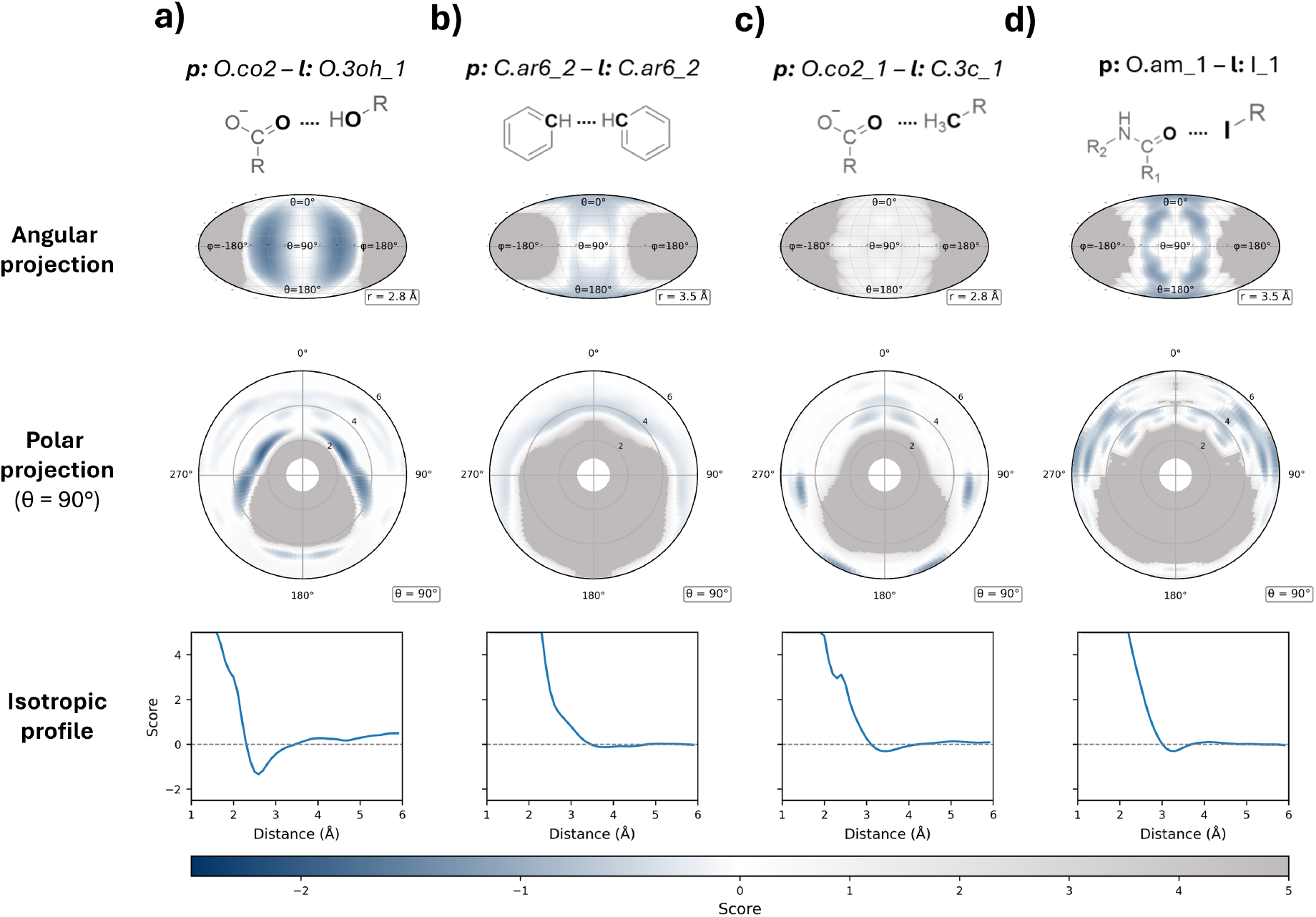
Anisotropic knowledge-based potential scores for four representative protein–ligand interaction types. Each column corresponds to a distinct interaction pair, identified by internal atom typing names and schematic depictions (top row): **a)** carboxylate O interacting with an alcohol O, **b)** unsubstituted, aromatic C interacting with another unsubstituted, aromatic C, **c)** carboxylate O interacting with a methyl C, and **d)** amide O interacting with I. The second row shows Mollweide projections of the score mapped onto the spherical surface at a fixed radial distance from the protein atom (*r* = 2.8 Å for **(a**,**d)**; *r* = 3.5 Å for **(b**,**c)**), revealing the directional preferences of each interaction. The third row displays polar plots of the score within the azimuthal plane (*θ* = 90 °), with the radial axis corresponding to the protein–ligand distance (Å). The bottom row shows the isotropic radial score profile from DESPOT-iso. Negative scores (blue) indicate geometries observed more frequently than expected from a reference distribution, while positive scores (gray) indicate disfavored geometries. The strong angular localization in **a)** and **b)** reflects the well-defined directionality of hydrogen bonding and *π*–*π* interactions, respectively.

Across all interaction types, DESPOT recovers a clear spectrum of directional specificity. Hydrogen-bonding interactions exhibit the strongest angular localization: favorable regions concentrate tightly along lone-pair directions, with well-defined preferences along the azimuthal axis *ϕ* while remaining comparatively uniform along *θ* (Figure 3a; Figure S3). Aromatic stacking interactions similarly display pronounced anisotropy, with favorable geometries concentrated perpendicular to the aromatic plane, consistent with face-to-face *π*–*π* stacking (Figure 3b; Figure S4). By contrast, interactions lacking a strong electronic driving force — such as alkyl contacts with polar groups — produce diffuse angular distributions dominated by steric exclusion rather than directional complementarity (Figure 3c). Between these extremes, interactions such as halogen contacts with amide oxygens show complex directional patterns of intermediate strength (Figure 3d; Figure S5).

Taken together, these profiles demonstrate that DESPOT captures physically meaningful interaction patterns without any explicit parametrization of interaction geometry. The directional features they reveal cannot be represented by radial distance profiles alone, underscoring the importance of incorporating angular information into knowledge-based scoring. Compared to the large residue-level reference frames of KORP-PL, DESPOT offers far higher interpretability at the level of individual interactions.

### Dual applications: pose scoring and MIF generation

A second advantage of DESPOT is that its inverted probabilistic formulation supports both pose scoring and MIF generation as two evaluations of the same underlying model. Figure 4 illustrates both applications for a biotin-bound protein.

**Figure 4:**
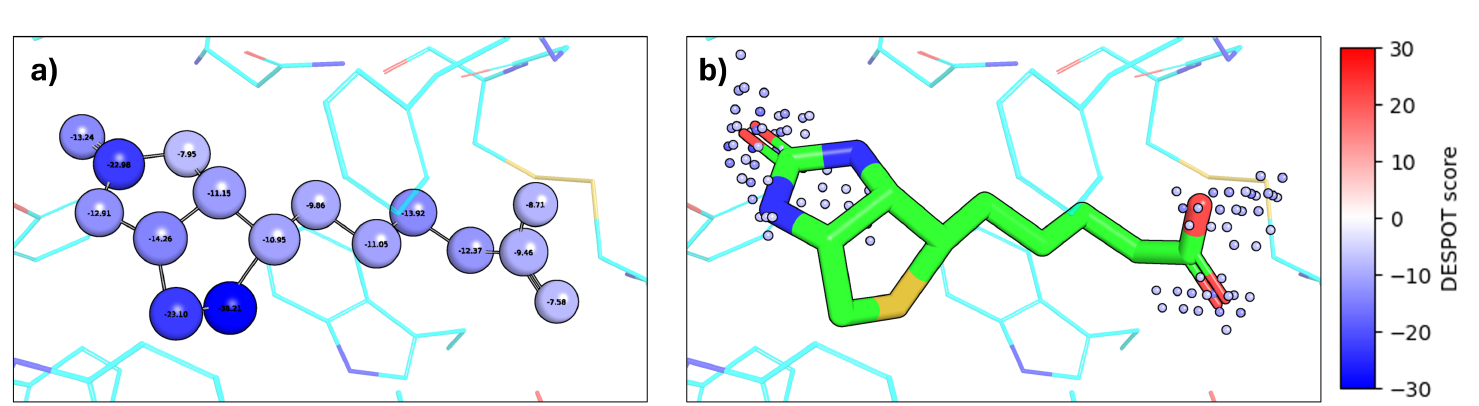
Dual applications of DESPOT: pose scoring and MIF generation. Illustration of DESPOT’s unified framework using a biotin-bound protein. **a)** Pose scoring: each ligand atom is colored by its DESPOT score, representing the sum of pairwise interactions with nearby protein atoms. Stronger negative scores (darker blue) indicate more favorable placement. **b)** Molecular interaction field generation: DESPOT potentials are evaluated on a regular three-dimensional grid without any ligand present, producing a multi-channel volumetric representation. Shown are all points in the carboxylate oxygen (*O*.*co*2_1) channel with negative values, with grid points colored by interaction score. The three pronounced energy minima cluster around the crystallographic positions of biotin’s sp2-hybridized oxygens, demonstrating that DESPOT MIFs encode chemically meaningful binding-site features.

In a pose scoring context, each ligand atom receives a score representing the sum of its pairwise interactions with nearby protein atoms, with negative scores indicating favorable placement. Because scores are resolved at the individual atom level, medicinal chemists can immediately identify which functional groups contribute favorably to binding and, crucially, which substituents occupy suboptimal positions and may benefit from further optimization. This per-atom decomposition moves beyond a single aggregate docking score to provide an interpretable diagnostic of ligand–protein complementarity that can directly inform lead optimization campaigns (Figure 4a).

In a MIF generation setting on the other hand, the same potentials are evaluated over a regular three-dimensional grid in the absence of any ligand, yielding a multi-channel representation in which each channel corresponds to a ligand atom type. In the example of Figure 4b, the carboxylate oxygen channel reveals three pronounced energy minima which cluster around the positions of biotin’s sp2-hybridized oxygens, demonstrating that DESPOT MIFs encode chemically meaningful binding-site features. MIFs thus complement per-atom pose scores by highlighting regions of unexploited binding-site potential where no ligand atom is currently positioned, offering a spatial guide for scaffold elaboration and fragment growing. These dense, multi-channel grids could also serve as natural inputs for future binding-site representation learning and structure-based design workflows [41, 42].

### CASF-2016 benchmark performance

DESPOT and its three variants — the isotropic DESPOT-iso, the residue-level DESPOT-screen and the combined score DESPOT-combo — were evaluated against established knowledge-based and empirical baselines (Figure 5).

**Figure 5:**
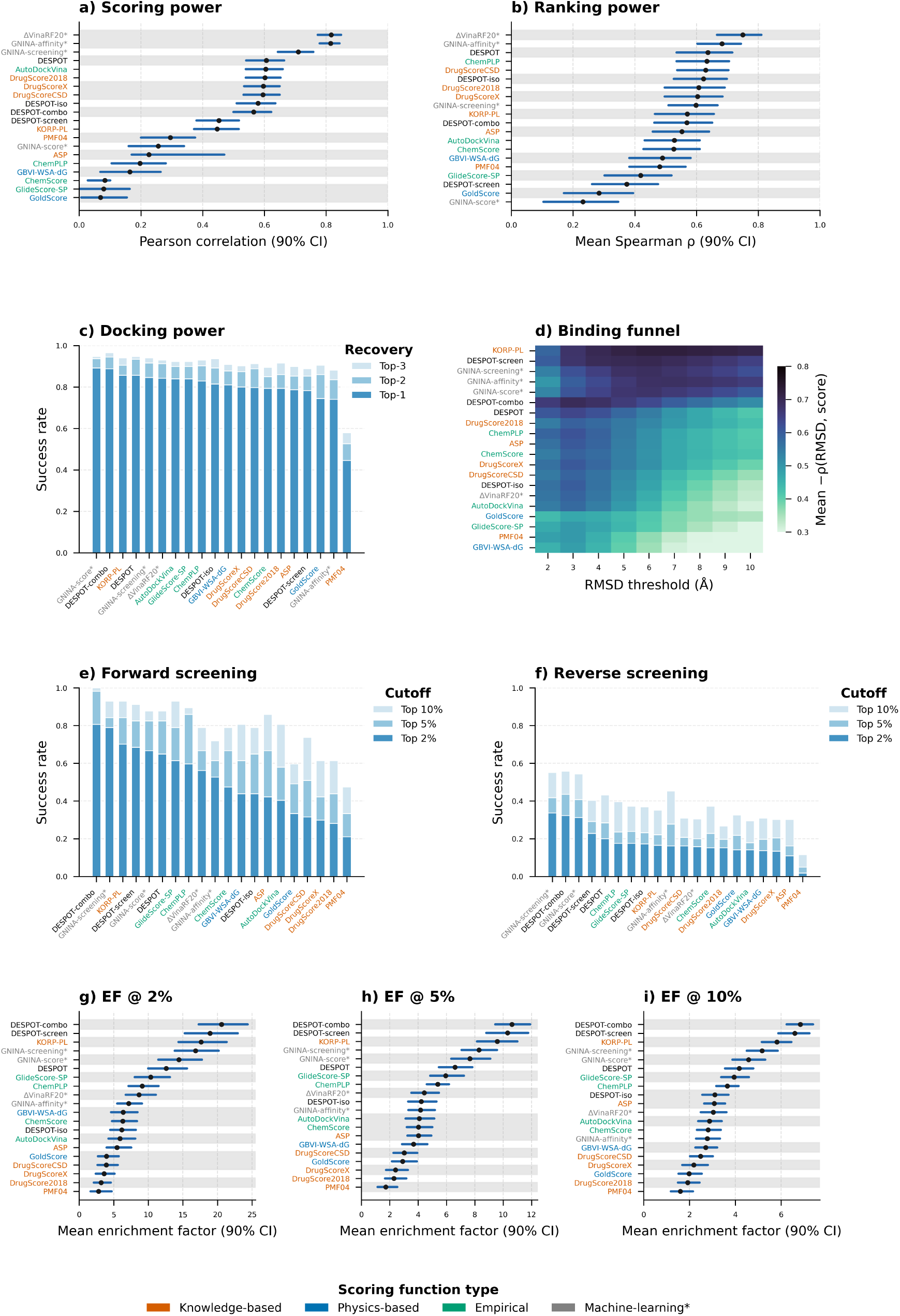
CASF-2016 benchmark performance. **a)** Scoring power: Pearson correlation between predicted scores and experimental binding affinities. **b)** Ranking power: mean Spearman correlation for ranking ligands within protein targets. Black dots indicate point estimates; blue bars indicate 90% bootstrap confidence intervals. Scoring functions on the same white/gray band are statistically indistinguishable (Friedman-Nemenyi test, p *>* 0.10). **c)** Docking power: success rate for identifying near-native poses (RMSD *<* 2.0 Å) at top-1, top-2, and top-3 rankings. **d)** Binding funnel analysis: Spearman correlation between score and RMSD as a function of RMSD threshold. **(e-f)** Screening power: success rate for forward screening (identifying correct ligands for targets) and reverse screening (identifying correct targets for ligands) at various ranking thresholds. **(g-i)** Enrichment factors at 2%, 5%, and 10% thresholds. The machine-learning potentials ΔVinaRF20 and GNINA are marked as a separate category, as there is significant bleed-through between their training data and the CASF-2016 core set.

#### Scoring and ranking power

The scoring power test measures the Pearson correlation between predicted scores and experimental affinities across 285 complexes (Figure 5a). DESPOT achieves a correlation of 0.610, competitive with established knowledge-based approaches such as DrugScore2018 and DrugScoreX. DESPOT and DESPOT-iso achieve nearly identical scoring power, indicating that anisotropy provides minimal advantage for affinity prediction of correctly positioned ligands. We attribute this to radial potentials already capturing the dominant statistical contributions to affinity (hydrophobic packing and effective entropic preferences), such that, for native poses, where directional constraints are satisfied by construction, the additional angular information has limited effect.

Ranking power parallels these findings (Figure 5b): DESPOT (Spearman’s *ρ* = 0.639) and the DrugScore variants all perform at comparable levels. By contrast, the residue-level DESPOT-screen performs poorly on both criteria (Pearson 0.430, Spearman 0.305).

#### Docking power

Docking power assesses the ability to identify near-native poses among decoys of the same ligand. DESPOT-combo achieves a top-1 success rate of 89.5% (255/285 targets), outperforming all other considered scoring functions except for GNINA-score in this challenge (Figure 5c). The binding-funnel analysis (Figure 5d) reveals a more surprising pattern: despite its modest top-3 pose recovery, DESPOT-screen maintains a strong rank correlation between score and RMSD even at very long ranges (up to 10 Å). These two observations seem to point in opposite directions — weak discrimination among the best poses, yet a well-behaved global funnel — and below we return to the coarse-graining mechanism that we believe reconciles them.

#### Screening power

Finally, the virtual screening setting reveals the largest differences among methods and provides the most compelling evidence for the value of anisotropy. In forward screening, DESPOT, DESPOT-screen, and DESPOT-combo achieve top-2% success rates of 38/57, 45/57, and 46/57, respectively, significantly outperforming DESPOT-iso (24/57) and all other isotropic KBPs (Figure 5e). Enrichment factors at the 2%, 5%, and 10% thresholds confirm this trend (Figure 5g-i): the three anisotropic models significantly outperform every isotropic KBP (*p* ≪ 0.0001 for all enrichment factors), as well as established empirical docking scores such as ChemPLP, GNINA, and ChemScore. DESPOT-screen also performs significantly better than its closest analog, KORP-PL — a gain we ascribe to (i) improved smoothing of the JPD and (ii) the greater quantity and quality of the training data.

Reverse screening is far more challenging: all tested scoring functions perform poorly, and only DESPOT-combo and GNINA achieve top-2% recovery above 30% (Figure 5f). Even so, the anisotropic KBPs are among the best-performing methods in this challenging setting, again clearly outperforming their isotropic variants. The weak overall performance likely reflects the difficulty of comparing docking scores across structurally diverse targets with different binding-site geometries and physicochemical properties; combining the atom-level DESPOT and residue-level DESPOT-screen scores nonetheless improves retrieval of the correct targets substantially.

#### A note of caution on machine-learning scoring functions

The CASF-2016 results of ΔVinaRF20 and GNINA are discussed separately here, as these ML-based scoring functions overlap substantially with CASF-2016 in both training data and training objectives [35, 39] (Figure 5). ΔVinaRF20 shows exceptional scoring and ranking power (Pearson: 0.816, Spearman: 0.750), but it was trained to predict affinities using PDBBind, and its train-test split excluded only the exact PDB IDs present in CASF-2016 — leaving substantial bleed-through from similar complexes spanning both sets. Under the 70%-sequence / 70%-ligand similarity clustering applied for DESPOT, a further 429 PDBBind entries are identified that should have been removed for a proper split.

GNINA was trained on CrossDocked2020, which again overlaps considerably with the CASF-2016 complexes [43]. Because its training objectives likewise include affinity prediction and near-native pose retrieval, its strong scoring (0.814), ranking (0.682), and docking power (89.5% top-1 recovery) are unsurprising.

#### Effects of training-set choice

The analyses above all use DESPOT models trained on CROWN. In our companion paper on CROWN [23], DESPOT and DESPOT-screen were also retrained using either the raw crystal structures from CROWN or HiQBind [21] as training resource. Training on CROWN rather than HiQBind improved both ranking and docking power: DESPOT’s mean Spearman correlation between score and affinity rose from 0.509 to 0.637, and DESPOT-screen’s from 0.235 to 0.374, while DESPOT-screen’s top-1 near-native pose recovery increased from 0.724 to 0.785. Differences between the energy-minimized and raw CROWN variants were much smaller and largely negligible for this benchmark. We refer to the companion paper [23] for the full comparison.

#### Summary of Benchmark Results

DESPOT, DESPOT-screen, and especially DESPOT-combo demonstrate consistent, competitive performance across all CASF-2016 criteria. While anisotropy yields little improvement in affinity prediction over classical DrugScore algorithms, it leads to substantial gains in pose discrimination and virtual screening. We hypothesize that this difference arises because misplaced docking poses violate directional preferences invisible to radial models, while native crystal poses already satisfy them, thus giving anisotropic methods a discriminative signal that radial ones lack.

This pattern — a small gap for near-native poses but a pronounced one when discriminating decoys mirrors observations from prospective blind challenges. In the SAMPL4 study, Voet et al. [44] found that expert “in cerebro” filtering of docking results boosted enrichment threefold over automated docking alone, with the experts’ advantage stemming from recognizing geometrically implausible interactions such as distorted hydrogen-bond angles or unrealistic aromatic stacking. DESPOT’s anisotropic potentials encode precisely this kind of directional knowledge in a fully automated manner, learned from over 110,000 curated, energy-minimized complexes. In this sense, DESPOT functions as encoded chemical intuition, automatically penalizing the geometric violations that experienced modelers recognize by eye but that distance-only methods cannot detect.

The different discretizations of DESPOT and DESPOT-screen shape performance in complementary ways: DESPOT’s atom-level potentials perform best in scoring, ranking, and docking power, whereas the residue-level DESPOT-screen is better in both forward and reverse screening. Returning to the funnel behavior noted above, we attribute DESPOT-screen’s combination of modest top-3 recovery and strong long-range correlation between score and RMSD to a single property of coarse-graining. Lacking atomic resolution, DESPOT-screen cannot discriminate finely among the closely spaced near-native poses that dominate the top rankings, which limits its top-3 recovery. The same coarse-graining, however, allows the receptor to make implicit conformation changes. CASF-2016 generates its docking decoys against a rigid crystal structure, so an atom-level model like DESPOT scores every pose against fixed side-chain rotamers, while DESPOT-screen is largely blind to them. Within this rigid-receptor benchmark, that blindness prevents DESPOT-screen from rewarding precise side-chain complementarity. The same insensitivity, however, should make the model more robust whenever the receptor conformation departs from the cognate crystal structure, as is likely in the long-range RMSD regime of docking and in the cross-docking regime of forward and reverse screening. DESPOT-screen’s poor scoring and ranking performance is harder to rationalize. The argument of Kadukova et al. [13], relating poor performance to the independence of energy-contribution terms, does not appear to apply here, since this independence holds for all KBP algorithms. DESPOT-combo, finally, unites the atom-level and residue-level models. Rather than averaging their performances, this simple combined score behaves as a consensus score: it retains strong scoring power while significantly improving docking power and both forward and reverse screening.

Taken together, these results show that DESPOT and DESPOT-screen are complementary. The fine-grained atom-level DESPOT excels at the exact placement of docked atoms, while the coarser DESPOT-screen implicitly captures flexibility: by ignoring exact side-chain rotamers it effectively tolerates the conformational adjustments a flexible receptor would make, behaving as though a degree of induced fit were built in. Combining the two levels of information yields more consistent and confident predictions.

### A unified probabilistic and spectral framework

Two aspects of DESPOT generalize beyond the present application. First, DrugScore, PMF, ASP, KORP-PL, and DESPOT can all be read as different conditionings and reference states of a single JPD *P* (*p, l*, **x**). Stating a potential in these terms makes explicit what it can and cannot express: most importantly, that only an occupancy-conditioned form is defined over empty space and can therefore represent steric exclusion and generate MIFs. We propose this as a useful organizing principle for the field: a knowledge-based potential is most cleanly specified and compared by which conditional and reference of *P* (*p, l*, **x**) it estimates.

Second, the spectral treatment of angular smoothing is, to our knowledge, new to knowledge-based potentials and transfers directly to any directional statistical potential. By smoothing intrinsically on the sphere rather than in real space, it decouples angular resolution from smoothing artifacts.

### Future directions

Although developed here for protein–small molecule interactions, the DESPOT framework is not inherently limited to this domain. The same directional preferences that govern hydrogen bonds and aromatic contacts in protein–ligand recognition also shape protein–protein and protein–nucleic acid interfaces. Extending DESPOT to these interaction types would require only a domain-specific training set of sufficient size and quality — a precedent established by DrugScore-PPI for isotropic protein–protein potentials [45]. Anisotropic potentials may prove especially valuable here, where larger, flatter interfaces amplify the role of directional complementarity relative to the buried-surface-area effects that dominate simple contact-based models.

A second direction is to use DESPOT’s anisotropic interaction grids as input representations for deep learning. The multi-channel MIFs encode rich, spatially resolved physicochemical information about binding sites in a ligand-independent manner, making them natural candidates for structure-based binding-site comparison and classification, where they could complement (or improve upon) approaches relying on sequence or residue-level features [42]. More broadly, integrating learned geometric interaction preferences into neural architectures could bridge the interpretability of knowledge-based methods with the representational flexibility of deep learning.

## Conclusion

We have presented DESPOT, a fine-grained, atom-level anisotropic knowledge-based scoring framework that extends classical distance-dependent potentials to full three-dimensional interaction models. A symmetry-aware geometric discretization preserves the simplicity and interpretability of classical KBPs while capturing the pronounced directional preferences that govern hydrogen bonding, aromatic stacking, halogen bonding, and other non-covalent interactions. Because knowledge-based potentials can be read as different conditionings of a single joint distribution *P* (*p, l, x*), DESPOT’s occupancy-conditioned, atom-level score has a natural complement: a geometry-conditioned, residue-level formulation (DESPOT-screen) that trades atomic resolution for an implicit tolerance of receptor flexibility. Summing the two inverse-Boltzmann scores yields DESPOT-combo, which behaves as a consensus rather than a compromise — retaining the strong scoring and ranking of the atom-level model while substantially improving docking power and both forward and reverse screening. Through its per-atom score decomposition, DESPOT supports interactive structure-based optimization workflows; through its occupancy-conditioned reference state, it unifies pose scoring and molecular interaction field generation within a single model. On CASF-2016, DESPOT-combo substantially outperforms established scoring functions in docking and screening power, with the advantage most pronounced when discriminating geometrically implausible poses. Our analyses confirm that data curation and rigorous train–test splitting are essential for deriving reliable anisotropic potentials and obtaining realistic performance estimates. DESPOT thus establishes an interpretable, data-driven foundation for direction-aware modeling of protein–ligand interactions, bridging the interpretability of knowledge-based potentials and the geometric expressiveness of physics-based force fields.

## Abbreviations

BCa: bias-corrected and accelerated (bootstrap)
CASF: Comparative Assessment of Scoring Functions
CCD: Chemical Component Dictionary
DH2: Driscoll-Healy (equiangular spherical grid)
ECFP4: extended-connectivity fingerprint (diameter 4)
EF: enrichment factor
JPD: joint probability distribution
KBP: knowledge-based potential
MIF: molecular interaction field
QED: quantitative estimate of drug-likeness
RMSD: root-mean-square deviation
VSEPR: valence shell electron pair repulsion

## Associated Content

### Supporting Information

(PDF). The full atom-typing scheme and refinement rules (Table S1); geometric definition of spherical voxels (Figure S1); atom type occurrence distributions of DESPOT (Figure S2); anisotropic potential profiles for hydrogen-bonding interactions with charged amine nitrogen (Figure S3), unsubstituted aromatic ring carbons (Figure S4), and halogen bonds with amide oxygen (Figure S5).

## Data and Software Availability

The complete CROWN dataset is freely available at https://crown.lbmd.be under a CC-BY 4.0 license. For each protein–ligand complex, protonated receptor structures are provided as PDB files and ligands with any associated cofactors as SDF files; both the corrected crystal coordinates prior to energy minimization and the final energy-minimized coordinates are included. Each entry is accompanied by crystallographic quality metrics (resolution, RSR, RSCC), biological annotations (UniProt ID, CATH classification, source organism), and ligand descriptors (SMILES, Murcko scaffold, molecular weight). The full dataset can be downloaded as a bulk archive at https://zenodo.org/records/20825315.

DESPOT is available as open-source software at https://github.com/KUL-LBMD/DESPOT under the MIT license. The repository includes scripts for training and inference with all DESPOT variants, along with documentation, environment specifications, and example usage. All pre-trained DESPOT models are available at https://zenodo.org/records/20829559.

## Author Contributions

R.P.: Conceptualization, Methodology, Data Curation, Investigation, Formal Analysis, Software, Writing – Original Draft. W.V.E. & A.S.: Validation, Writing – Review Editing. B.B. & A.A.: Methodology, Supervision, Writing – Review Editing. Y.M. & A.V.: Conceptualization, Supervision, Writing – Review & Editing.

## Acknowledgements

Robin Poelmans and Ahmed Shemy acknowledge support from the Research Foundation - Flanders (FWO) through PhD fellowships (Grant No. 1176225N and G095522N, respectively). Arnout Voet and Bence Bruncsics acknowledge funding through the Citribel Research Chair in Protein Engineering for Circular Waste Processes. Adam Arany and Yves Moreau are affiliated with Leuven.AI and received funding from the Flemish Government (AI Research Program).

