## Supplementary Information for "DESPOT: Direction-Enhanced Scoring POTentials for Scoring of Protein-Ligand Interactions"

#### Contents

**Supporting Information** (PDF). The full atom-typing scheme and refinement rules (Table S1); geometric definition of spherical voxels (Figure S1); atom type occurrence distributions of DESPOT (Figure S2); anisotropic potential profiles for hydrogen-bonding interactions with charged amine nitrogen (Figure S3), unsubstituted aromatic ring carbons (Figure S4), and halogen bonds with amide oxygen (Figure S5).

### Supporting Information

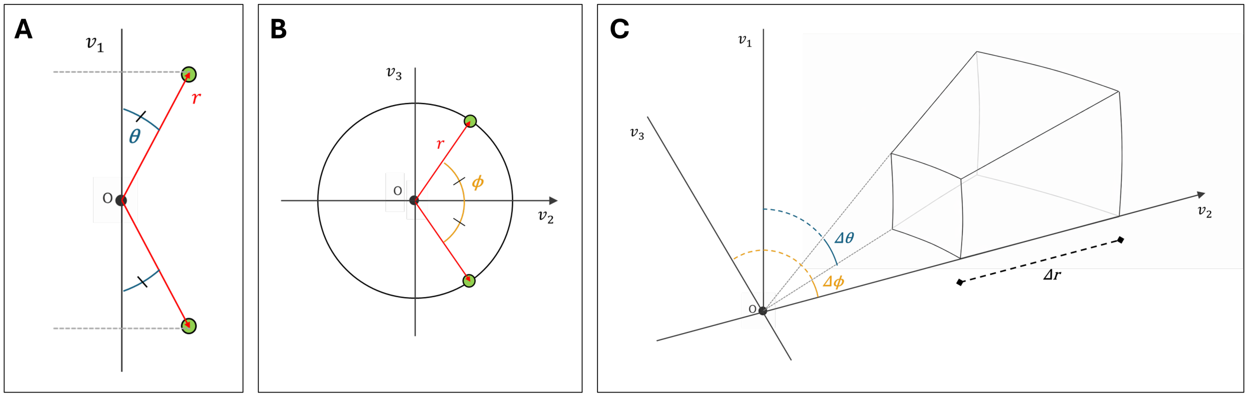


Figure S1: **Geometric definition of spherical voxels used in DESPOT.**
**(A)** Treatment of $v_1$ as an unsigned axis: because the plane normal can point in either direction, the polar angle $\theta$ is restricted to [0°–90°] with mirror symmetry about the plane. **(B)** Treatment of $v_3$ as an unsigned axis: the inherited directional ambiguity restricts the azimuthal angle $\phi$ to [0°–180°]. **(C)** Resulting spherical voxel geometry for fully anisotropic atoms, discretized in $(r, \theta, \phi)$ with $\Delta r = 0.1$ Å and $\Delta\theta = \Delta\phi = 3.0°$. Each voxel captures interactions from four equivalent regions of space due to the unsigned axis treatment.

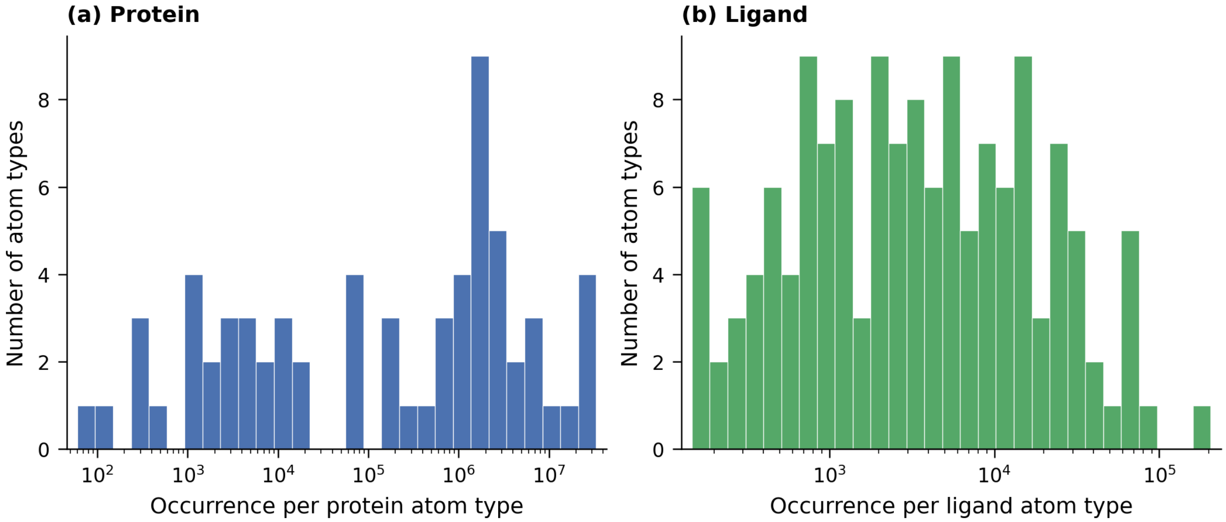


Figure S2: **Atom type occurrence distributions of DESPOT.**
**a)** Occurrence distribution of protein atom types used by DESPOT. **b)** Occurrence distribution of ligand atom types used by DESPOT. x-axis displayed in log-scale for both plots.

Table S1: Atom-type definitions used in this work. 'In data set' indicates whether each atom type is assigned to the protein and/or ligand side (✓), which is independent of its raw occurrence counts $N_{\text{prot}}$ and $N_{\text{lig}}$. #H denotes the number of attached hydrogens; a slash (/) marks attributes that do not apply. Ref. frame is the local reference frame (anisotropic, axial, or isotropic).

| | In data set | | | | | | | | |
|---|---|---|---|---|---|---|---|---|---|
| **Atom type** | **Prot.** | **Lig.** | $N_{\text{prot}}$ | $N_{\text{lig}}$ | **Elem.** | **Hybrid.** | **#H** | **Ref. frame** | **Definition** |
| O.am_1 | ✓ | ✓ | 33,338,575 | 39,623 | O | sp2 | 0 | Anisotropic | Amide O |
| C.am_3 | ✓ | ✓ | 33,318,170 | 28,213 | C | sp2 | 0 | Axial | Carbonyl C in amide |
| N.am_2 | ✓ | ✓ | 29,361,447 | 26,375 | N | sp2 | 1 | Anisotropic | Secondary amide N |
| C.3n_3 | ✓ | ✓ | 27,190,576 | 13,870 | C | sp3 | 1 | Axial | sp3 C bound to N + 2 other heavy atoms |
| C.3c_2 | ✓ | ✓ | 26,657,559 | 83,371 | C | sp3 | 2 | Anisotropic | Secondary C (bound to C only) |
| C.3c_1 | ✓ | ✓ | 17,762,383 | 63,650 | C | sp3 | 3 | Axial | Methyl group (bound to C only) |
| C.ar6_2 | ✓ | ✓ | 13,164,477 | 204,787 | C | sp2 | 1 | Anisotropic | Unsubstituted C in aromatic 6-ring (C-only ring) |
| O.co2_1 | ✓ | ✓ | 7,579,503 | 45,132 | O | sp2 | 0 | Anisotropic | Carboxylate O |
| C.3c_3 | ✓ | ✓ | 6,801,518 | 7,267 | C | sp3 | 1 | Axial | Tertiary C (bound to C only) |
| C.3n_2 | ✓ | ✓ | 5,508,828 | 28,329 | C | sp3 | 2 | Anisotropic | sp3 C bound to N + 1 other heavy atom |
| C.co2_3 | ✓ | ✓ | 3,789,663 | 22,566 | C | sp2 | 0 | Axial | Carboxylate C |
| O.3oh_1 | ✓ | ✓ | 3,679,111 | 75,114 | O | sp3 | 1 | Axial | Alcohol O |
| N.gu_1 | ✓ | ✓ | 3,101,557 | 5,192 | N | sp2 | [1,2] | Anisotropic | Guanidine N with 1 heavy neighbor |
| C.ar6_3 | ✓ | ✓ | 2,963,312 | 65,420 | C | sp2 | 0 | Axial | Substituted C in aromatic 6-ring (C-only ring) |
| C.3r5_2 | ✓ | ✓ | 3,003,869 | 13,925 | C | sp3 | 2 | Anisotropic | sp3 C in 5-membered ring |
| N.am_1 | ✓ | ✓ | 2,464,532 | 1,897 | N | sp2 | 2 | Anisotropic | Primary amide N |
| C.ar5_2 | ✓ | ✓ | 2,162,562 | 23,847 | C | sp2 | 1 | Anisotropic | Unsubstituted C in aromatic 5-ring |
| C.oh_2 | ✓ | ✓ | 1,877,955 | 14,991 | C | sp3 | 2 | Anisotropic | Primary alcohol |
| N.4_1 | ✓ | ✓ | 1,796,438 | 8,043 | N | sp3 | 3 | Axial | Primary amine, charged |
| C.3r5x_3 | ✓ | ✓ | 1,779,792 | 24,632 | C | sp3 | 1 | Axial | sp3 C in 5-membered ring with hetero-atom neighbor |
| C.oh_3 | ✓ | ✓ | 1,721,680 | 13,236 | C | sp3 | 1 | Axial | Secondary alcohol |
| C.ar6x_3 | ✓ | ✓ | 1,635,303 | 70,586 | C | sp2 | 0 | Axial | C in aromatic 6-ring with heteroatom substituent |
| N.gu_2 | ✓ | ✓ | 1,584,192 | 14,596 | N | sp2 | [0,1] | Anisotropic | Guanidine N with 2 heavy neighbors |
| N.am_3 | ✓ | ✓ | 1,537,308 | 10,437 | N | sp2 | 0 | Axial | Tertiary amide N |
| C.guh_3 | ✓ | ✓ | 1,540,231 | 732 | C | sp2 | 0 | Axial | Guanidinium C |
| C.3r5x_2 | ✓ | ✓ | 1,499,975 | 7,858 | C | sp3 | 2 | Anisotropic | sp3 C in 5-membered ring with hetero-atom neighbor |
| C.ar5_3 | ✓ | ✓ | 1,339,357 | 34,229 | C | sp2 | 0 | Axial | Substituted C in aromatic 5-ring |
| N.ar5p_2 | ✓ | ✓ | 1,306,554 | 5,586 | N | sp2 | 1 | Anisotropic | Unsubstituted, charged N in 5-membered aromatic ring |
| C.3s_2 | ✓ | ✓ | 1,232,298 | 3,490 | C | sp3 | 2 | Anisotropic | sp3 C coupled to S |
| O.ph_1 | ✓ | ✓ | 1,135,436 | 11,804 | O | sp2 | 1 | Anisotropic | Phenol O |
| N.ar5_2 | ✓ | ✓ | 846,286 | 8,267 | N | sp2 | 0 | Anisotropic | Unsubstituted, uncharged N in 5-membered aromatic ring |
| S.3_2 | ✓ | ✓ | 751,758 | 2,506 | S | sp3 | 0 | Anisotropic | sp3 S coupled to 2 C's |
| C.3s_1 | ✓ | ✓ | 749,293 | 528 | C | sp3 | 3 | Axial | sp3 C coupled to S |
| S.csh_1 | ✓ | ✓ | 392,253 | 1,114 | S | sp3 | 1 | Axial | Thiol S |
| O.h2o_0 | ✓ | – | 322,181 | 0 | O | sp3 | 2 | Isotropic | Water |

*continued on next page*

| Atom type | In data set Prot. | In data set Lig. | $N_{\mathbf{prot}}$ | $N_{\mathbf{lig}}$ | Elem. | Hybrid. | #H | Ref. frame | Definition |
|---|---|---|---|---|---|---|---|---|---|
| C.ar6o_3 | ✓ | ✓ | 167,791 | 52,243 | C | sp2 | 0 | Axial | Substituted C in aromatic 6-ring (ortho relative to hetero-atom) |
| O.2p_1 | ✓ | ✓ | 156,749 | 33,980 | O | sp2 | 0 | Axial | Primary phosphate O |
| O.3p_2 | ✓ | ✓ | 143,954 | 12,328 | O | sp3 | 0 | Anisotropic | Connecting O between phosphates |
| C.et_2 | ✓ | ✓ | 76,183 | 16,366 | C | sp3 | 2 | Anisotropic | Ether C |
| O.3ret_2 | ✓ | ✓ | 72,936 | 18,292 | O | sp3 | 0 | Anisotropic | Ether O in ring |
| S.s_2 | ✓ | – | 87,017 | 34 | S | sp3 | 0 | Anisotropic | Disulphide bridges |
| P.o4_4 | ✓ | ✓ | 75,451 | 10,731 | P | sp3 | 0 | Isotropic | P coupled to 4 O's |
| C.3r6x_3 | ✓ | ✓ | 11,658 | 66,853 | C | sp3 | 1 | Axial | sp3 C in 6-membered ring with hetero-atom neighbor |
| C.ar6o_2 | – | ✓ | 52,253 | 20,319 | C | sp2 | 1 | Anisotropic | Unsubstituted C in aromatic 6-ring (ortho relative to hetero-atom) |
| C.ar6m_3 | – | ✓ | 44,990 | 23,037 | C | sp2 | 0 | Axial | Substituted C in aromatic 6-ring (meta relative to hetero-atom) |
| N.ar6_2 | – | ✓ | 43,082 | 22,523 | N | sp2 | 0 | Anisotropic | Unsubstituted, uncharged N in 6-membered aromatic ring |
| O.ke_1 | – | ✓ | 36,541 | 12,136 | O | sp2 | 0 | Anisotropic | Ketone O |
| N.ar5p_3 | – | ✓ | 41,207 | 6,149 | N | sp2 | 0 | Axial | Substituted, charged N in 5-membered aromatic ring |
| C.ar6m_2 | – | ✓ | 30,380 | 14,899 | C | sp2 | 1 | Anisotropic | Unsubstituted C in aromatic 6-ring (meta relative to hetero-atom) |
| N.mih_1 | – | ✓ | 36,229 | 6,920 | N | sp2 | [1,2] | Anisotropic | Primary amidine N |
| C.3r6_2 | – | ✓ | 6,942 | 35,619 | C | sp3 | 2 | Anisotropic | sp3 C in 6-membered ring |
| C.2r6x_3 | – | ✓ | 31,532 | 8,945 | C | sp2 | 0 | Axial | Non-aromatic sp2 C in 6-membered ring with hetero-atom neighbor |
| C.2c_2 | ✓ | ✓ | 19,362 | 14,925 | C | sp2 | 1 | Anisotropic | Secondary alkene |
| C.3r6x_2 | – | ✓ | 2,563 | 24,976 | C | sp3 | 2 | Anisotropic | sp3 C in 6-membered ring with hetero-atom neighbor |
| F_1 | – | ✓ | 1,716 | 18,227 | F | sp3 | 0 | Axial | Fluoride group |
| N.r6_3 | – | ✓ | 15,919 | 3,943 | N | sp3 | 0 | Axial | sp3 N in 6-membered ring |
| C.2r6_2 | – | ✓ | 16,323 | 2,512 | C | sp2 | 1 | Anisotropic | Non-aromatic sp2 C in 6-membered ring |
| C.es_3 | ✓ | ✓ | 15,076 | 3,309 | C | sp2 | 0 | Axial | Ester C |
| O.3n_2 | – | ✓ | 17,050 | 311 | O | sp3 | 0 | Anisotropic | O coupled to N and C |
| N.im_2 | – | ✓ | 13,989 | 2,971 | N | sp2 | 1 | Anisotropic | Secondary imide N |
| O.3eta_2 | – | ✓ | 1,724 | 14,375 | O | sp3 | 0 | Anisotropic | Ether O with aromatic ring as substituent |
| O.es_1 | ✓ | ✓ | 11,533 | 3,986 | O | sp2 | 0 | Anisotropic | Carbonyl O in ester |
| O.3es_2 | ✓ | ✓ | 8,066 | 5,850 | O | sp3 | 0 | Anisotropic | Ester alkoxy oxygen |
| C.3r6_3 | – | ✓ | 2,629 | 11,060 | C | sp3 | 1 | Axial | sp3 C in 6-membered ring |
| C.2c_1 | ✓ | ✓ | 11,254 | 1,392 | C | sp2 | 2 | Anisotropic | Primary alkene |
| C.3n_1 | – | ✓ | 1,733 | 10,881 | C | sp3 | 3 | Axial | Methyl C coupled to N |
| N.4_2 | – | ✓ | 5,941 | 4,823 | N | sp3 | 2 | Anisotropic | Secondary amine, charged |
| O.3et_2 | ✓ | ✓ | 2,078 | 8,411 | O | sp3 | 0 | Anisotropic | Ether oxygen |
| Cl_1 | – | ✓ | 907 | 9,163 | Cl | sp3 | 0 | Axial | Chloride group |
| C.1_2 | ✓ | ✓ | 6,563 | 3,453 | C | sp | 0 | Isotropic | Secondary alkyne |
| C.et_1 | – | ✓ | 1,016 | 8,905 | C | sp3 | 3 | Axial | Ether C |

*continued on next page*

Table S1 – *continued from previous page*

| Atom type | In data set Prot. | Lig. | $N_{prot}$ | $N_{lig}$ | Elem. | Hybrid. | #H | Ref. frame | Definition |
|---|---|---|---|---|---|---|---|---|---|
| N.ar5n_2 | – | ✓ | 756 | 8,972 | N | sp2 | 0 | Anisotropic | Unsubstituted, uncharged N in 5-membered aromatic ring next to N |
| C.ar6p_3 | – | ✓ | 792 | 7,359 | C | sp2 | 0 | Axial | Substituted C in aromatic 6-ring (para relative to hetero-atom) |
| C.2r6_3 | – | ✓ | 1,632 | 6,179 | C | sp2 | 0 | Axial | Non-aromatic sp2 C in 6-membered ring |
| C.ar6p_2 | – | ✓ | 1,917 | 5,636 | C | sp2 | 1 | Anisotropic | Unsubstituted C in aromatic 6-ring (para relative to hetero-atom) |
| N.mih_2 | – | ✓ | 876 | 6,403 | N | sp2 | [0,1] | Anisotropic | Secondary amidine N |
| O.2s_1 | – | ✓ | 2,129 | 4,409 | O | sp2 | 0 | Axial | Primary sulphate O |
| C.2r5x_3 | ✓ | ✓ | 1,849 | 4,651 | C | sp2 | 0 | Axial | Non-aromatic sp2 C in 5-membered ring with hetero-atom neighbor |
| C.3r6_4 | – | ✓ | 1,427 | 4,960 | C | sp3 | 0 | Isotropic | sp3 C in 6-membered ring |
| N.4_3 | – | ✓ | 1,187 | 4,981 | N | sp3 | 1 | Axial | Tertiary amine, charged |
| C.2n_2 | ✓ | ✓ | 4,944 | 708 | C | sp2 | 1 | Anisotropic | Non-aromatic sp2 C bound to N and 1 other heavy atom |
| C.3r5_3 | – | ✓ | 884 | 4,712 | C | sp3 | 1 | Axial | sp3 C in 5-membered ring |
| N.gu_3 | ✓ | ✓ | 4,091 | 1,427 | N | sp2 | 0 | Axial | Guanidine N with 3 heavy neighbors |
| S.ar_2 | – | ✓ | 601 | 4,912 | S | sp2 | 0 | Anisotropic | S in aromatic ring |
| N.ar6p_3 | – | ✓ | 2,603 | 2,030 | N | sp2 | 0 | Axial | Substituted, charged N in 6-membered aromatic ring |
| C.2c_3 | – | ✓ | 946 | 3,585 | C | sp2 | 0 | Axial | Tertiary alkene |
| Zn | ✓ | – | 4,290 | 0 | Zn | / | / | Isotropic | Zn ion |
| C.o_3 | – | ✓ | 756 | 3,440 | C | sp2 | 0 | Axial | C of keto group |
| C.3r3_2 | – | ✓ | 334 | 3,737 | C | sp3 | 2 | Anisotropic | sp3 C in 3-membered ring |
| N.r6_2 | – | ✓ | 943 | 2,857 | N | sp3 | 1 | Anisotropic | sp3 N in 6-membered ring |
| C.et_3 | – | ✓ | 1,535 | 2,189 | C | sp3 | 1 | Axial | Ether C |
| N.3c_1 | ✓ | – | 3,603 | 94 | N | sp3 | 2 | Axial | Primary amine, uncharged |
| N.ar5np_3 | – | ✓ | 229 | 3,424 | N | sp2 | 0 | Axial | Substituted, charged N in 5-membered aromatic ring next to N |
| O.2n_1 | – | ✓ | 579 | 2,794 | O | sp2 | 0 | Anisotropic | Nitro O |
| C.3r6x_4 | – | ✓ | 332 | 2,896 | C | sp3 | 0 | Isotropic | sp3 C in 6-membered ring with hetero-atom neighbor |
| Mg | ✓ | – | 3,034 | 0 | Mg | / | / | Isotropic | Mg ion |
| C.3hal_4 | – | ✓ | 259 | 2,730 | C | sp3 | 0 | Isotropic | sp3 C bound to halogen |
| C.2r5x_2 | ✓ | – | 2,675 | 53 | C | sp2 | 1 | Anisotropic | Non-aromatic sp2 C in 5-membered ring with hetero-atom neighbor |
| N.aa_2 | – | ✓ | 322 | 2,389 | N | sp2 | [0,1] | Anisotropic | Secondary amine as aromatic substituent |
| P.o3_4 | – | ✓ | 762 | 1,804 | P | sp3 | 0 | Isotropic | P coupled to 3 O's and 1 C |
| O.ar_2 | – | ✓ | 295 | 2,188 | O | sp2 | 0 | Anisotropic | O in aromatic ring |
| N.1_1 | – | ✓ | 470 | 1,867 | N | sp | 0 | Axial | Nitrile N |
| N.oh_2 | – | ✓ | 1,004 | 1,327 | N | sp2 | 0 | Anisotropic | Nitroso group with 1 C neighbour |
| Br_1 | – | ✓ | 331 | 1,896 | Br | sp3 | 0 | Axial | Bromide group |

*continued on next page*

| Atom type | In data set Prot. | In data set Lig. | $N_{\mathbf{prot}}$ | $N_{\mathbf{lig}}$ | Elem. | Hybrid. | #H | Ref. frame | Definition |
|---|---|---|---|---|---|---|---|---|---|
| C.3r5x_4 | – | ✓ | 223 | 1,958 | C | sp3 | 0 | Isotropic | sp3 C in 5-membered ring with hetero-atom neighbor |
| N.ar5np_2 | – | ✓ | 177 | 1,930 | N | sp2 | 1 | Anisotropic | Unsubstituted, charged N in 5-membered aromatic ring next to N |
| N.r5_2 | ✓ | ✓ | 1,444 | 458 | N | sp3 | 1 | Anisotropic | sp3 N in 5-membered ring |
| C.1_1 | – | ✓ | 1,673 | 220 | C | sp | 1 | Axial | Primary alkyne |
| S.o3n0_4 | – | ✓ | 635 | 1,238 | S | sp3 | 0 | Isotropic | sp3 S coupled to O's and 1 C |
| N.3c_2 | – | ✓ | 1,513 | 234 | N | sp3 | 1 | Anisotropic | Secondary amine, uncharged |
| C.2r6x_2 | – | ✓ | 1,309 | 381 | C | sp2 | 1 | Anisotropic | Non-aromatic sp2 C in 6-membered ring with hetero-atom neighbor |
| C.3c_4 | – | ✓ | 303 | 1,364 | C | sp3 | 0 | Isotropic | Quaternary C (bound to C only) |
| N.o2_3 | – | ✓ | 256 | 1,321 | N | sp2 | 0 | Axial | Nitro N |
| C.3r3_3 | – | ✓ | 128 | 1,435 | C | sp3 | 1 | Axial | sp3 C in 3-membered ring |
| N.2c_2 | – | ✓ | 469 | 1,048 | N | sp2 | [0,1] | Anisotropic | sp2 N, secondary |
| S.2c_1 | ✓ | ✓ | 1,049 | 416 | S | sp2 | 0 | Anisotropic | sp2 S coupled to C |
| Mn | ✓ | – | 1,427 | 0 | Mn | / | / | Isotropic | Mn ion |
| N.aa_1 | – | ✓ | 168 | 1,186 | N | sp2 | [1,2] | Anisotropic | Primary amine as aromatic substituent |
| C.o_2 | ✓ | ✓ | 478 | 874 | C | sp2 | 1 | Anisotropic | C of terminal aldehyde |
| C.2n_3 | – | ✓ | 234 | 1,091 | C | sp2 | 0 | Axial | Non-aromatic sp2 C bound to N and 2 other heavy atoms |
| C.oh_4 | – | ✓ | 142 | 1,120 | C | sp3 | 0 | Isotropic | Tertiary alcohol |
| C.3r3x_3 | – | ✓ | 588 | 664 | C | sp3 | 1 | Axial | sp3 C in 3-membered ring with hetero-atom neighbor |
| N.r5_3 | – | ✓ | 443 | 746 | N | sp3 | 0 | Axial | sp3 N in 5-membered ring |
| Ca | ✓ | – | 1,155 | 0 | Ca | / | / | Isotropic | Ca ion |
| C.et_4 | – | ✓ | 450 | 673 | C | sp3 | 0 | Isotropic | Ether C |
| O.al_1 | ✓ | ✓ | 317 | 806 | O | sp2 | 0 | Anisotropic | Aldehyde O |
| C.3r4x_2 | – | ✓ | 111 | 988 | C | sp3 | 2 | Anisotropic | sp3 C in 4-membered ring with hetero-atom neighbor |
| S.r_2 | – | ✓ | 77 | 997 | S | sp3 | 0 | Anisotropic | sp3 S in ring |
| C.3r4_2 | – | ✓ | 120 | 944 | C | sp3 | 2 | Anisotropic | sp3 C in 3-membered ring |
| C.2r5_3 | – | ✓ | 95 | 884 | C | sp2 | 0 | Axial | Non-aromatic sp2 C in 5-membered ring |
| N.ar6n_2 | – | ✓ | 53 | 879 | N | sp2 | 0 | Anisotropic | Unsubstituted, uncharged N in 6-membered aromatic ring next to N |
| N.aa_3 | – | ✓ | 127 | 693 | N | sp2 | 0 | Axial | Tertiary amine as aromatic substituent |
| N.im_3 | – | ✓ | 101 | 709 | N | sp2 | 0 | Axial | Tertiary imide N |
| C.3p_2 | – | ✓ | 131 | 612 | C | sp3 | 2 | Anisotropic | sp3 C coupled to P and C |
| N.ar6np_3 | – | ✓ | 26 | 683 | N | sp2 | 0 | Axial | Substituted, charged N in 6-membered aromatic ring next to N |
| N.mih_3 | – | ✓ | 101 | 605 | N | sp2 | 0 | Axial | Tertiary amidine N |
| C.3r4x_3 | – | ✓ | 237 | 433 | C | sp3 | 1 | Axial | sp3 C in 4-membered ring with hetero-atom neighbor |

*continued on next page*

Table S1 – *continued from previous page*

| Atom type | In data set Prot. | Lig. | $N_{\mathbf{prot}}$ | $N_{\mathbf{lig}}$ | Elem. | Hybrid. | #H | Ref. frame | Definition |
|---|---|---|---|---|---|---|---|---|---|
| C.3r3x_4 | – | ✓ | 274 | 358 | C | sp3 | 0 | Isotropic | sp3 C in 3-membered ring with hetero-atom neighbor |
| N.4_4 | – | ✓ | 230 | 401 | N | sp3 | 0 | Isotropic | Quaternary amine, charged |
| C.3r5_4 | – | ✓ | 42 | 555 | C | sp3 | 0 | Isotropic | sp3 C in 5-membered ring |
| C.3n_4 | – | ✓ | 73 | 522 | C | sp3 | 0 | Isotropic | sp3 C bound to N + 3 other heavy atoms |
| N.2n_2 | – | ✓ | 94 | 449 | N | sp2 | [0,1] | Anisotropic | sp2 bound to N and C |
| I_1 | – | ✓ | 22 | 495 | I | sp3 | 0 | Axial | Iodine group |
| C.2r5_2 | – | ✓ | 131 | 263 | C | sp2 | 1 | Anisotropic | Non-aromatic sp2 C in 5-membered ring |
| C.3r4_3 | – | ✓ | 60 | 325 | C | sp3 | 1 | Axial | sp3 C in 3-membered ring |
| N.ar6p_2 | – | ✓ | 89 | 292 | N | sp2 | 1 | Anisotropic | Unsubstituted, charged N in 6-membered aromatic ring |
| N.3p_2 | – | ✓ | 145 | 147 | N | sp3 | 1 | Anisotropic | sp3 N bound to P |
| C.3hal_3 | – | ✓ | 23 | 269 | C | sp3 | 1 | Axial | sp3 C bound to halogen |
| C.am_2 | – | ✓ | 112 | 151 | C | sp2 | 1 | Anisotropic | Carbonyl C in terminal amide |
| Na | ✓ | – | 258 | 0 | Na | / | / | Isotropic | Na ion |
| O.2o_1 | ✓ | – | 245 | 0 | O | sp2 | 0 | Axial | Oxygen |
| O.3s_2 | – | ✓ | 76 | 167 | O | sp3 | 0 | Anisotropic | Connecting O between sulphates |
| S.o4n0_4 | – | ✓ | 54 | 170 | S | sp3 | 0 | Isotropic | Sulphate S |
| C.3hal_2 | – | ✓ | 32 | 185 | C | sp3 | 2 | Anisotropic | sp3 C bound to halogen |
| C.3s_3 | – | ✓ | 12 | 183 | C | sp3 | 1 | Axial | sp3 C coupled to S |
| K | ✓ | – | 104 | 0 | K | / | / | Isotropic | K ion |
| Cd | ✓ | – | 60 | 0 | Cd | / | / | Isotropic | Cd ion |

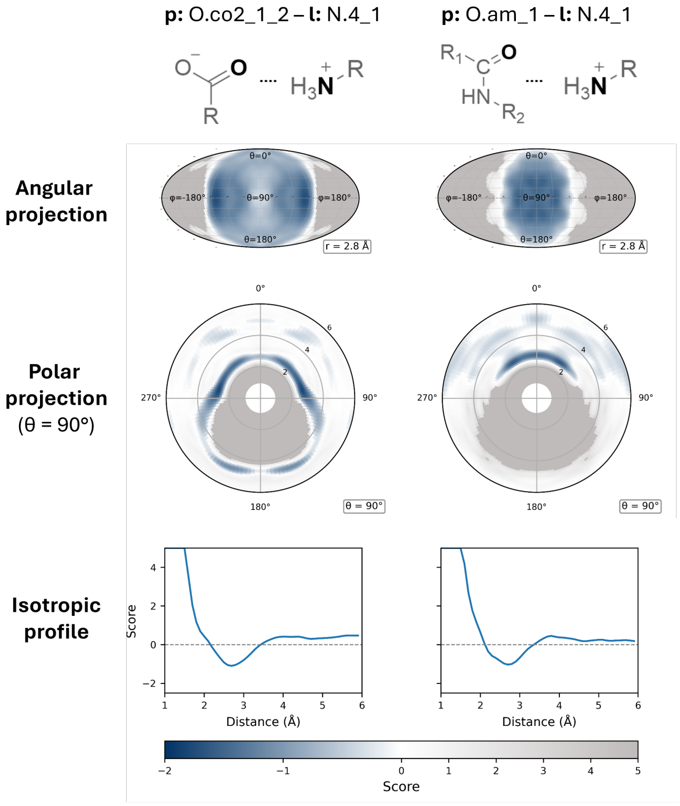


Figure S3: **Anisotropic knowledge-based potential scores for two hydrogen-bonding interaction types involving a charged amine nitrogen**. Each column corresponds to a distinct interaction pair, identified by internal atom typing names and schematic depictions (top row): carboxylate O interacting with a primary charged amine N and amide O interacting with a primary charged amine N. The second row shows Mollweide projections of the score mapped onto the spherical surface at a fixed radial distance from the protein atom ($r = 2.8$ Å for all columns), revealing the directional preferences of each interaction. The third row displays polar plots of the score within the azimuthal plane ($\theta = 90\ °$), with the radial axis corresponding to the protein–ligand distance (Å). The bottom row shows the isotropic radial score profile from DESPOT-iso. Negative scores (blue) indicate geometries observed more frequently than expected from a reference distribution, while positive scores (gray) indicate disfavored geometries. All three oxygen types show pronounced angular localization of favorable contacts along lone-pair directions, with subtle differences in the breadth and symmetry of the preferred regions reflecting the distinct electronic environments of the carboxylate, amide, and ketone carbonyl.

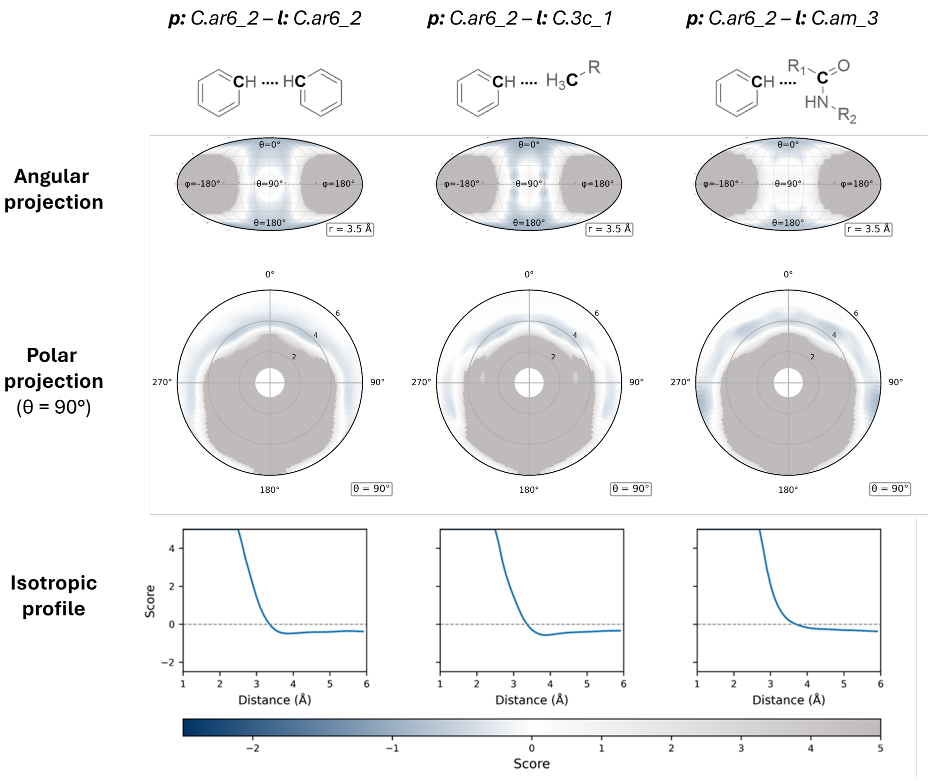


Figure S4: **Anisotropic knowledge-based potential scores for three interaction types involving an unsubstituted aromatic ring carbon.** Each column corresponds to a distinct interaction pair, identified by internal atom typing names and schematic depictions (top row): unsubstituted aromatic C interacting with another unsubstituted aromatic C, unsubstituted aromatic C interacting with a methyl C, and unsubstituted aromatic C interacting with an amide C. The second row shows Mollweide projections of the score mapped onto the spherical surface at a fixed radial distance from the protein atom (r = 3.5 Å for all columns), revealing the directional preferences of each interaction. The third row displays polar plots of the score within the azimuthal plane ($\theta = 90$ °), with the radial axis corresponding to the protein–ligand distance (Å). The bottom row shows the isotropic radial score profile from DESPOT-iso. Negative scores (blue) indicate geometries observed more frequently than expected from a reference distribution, while positive scores (gray) indicate disfavored geometries. The aromatic–aromatic interaction (column 1) displays strong angular localization along the axis perpendicular to the ring plane, consistent with face-to-face $\pi-\pi$ stacking, while the methyl and amide carbon interactions show progressively more diffuse angular distributions reflecting weaker directional driving forces.

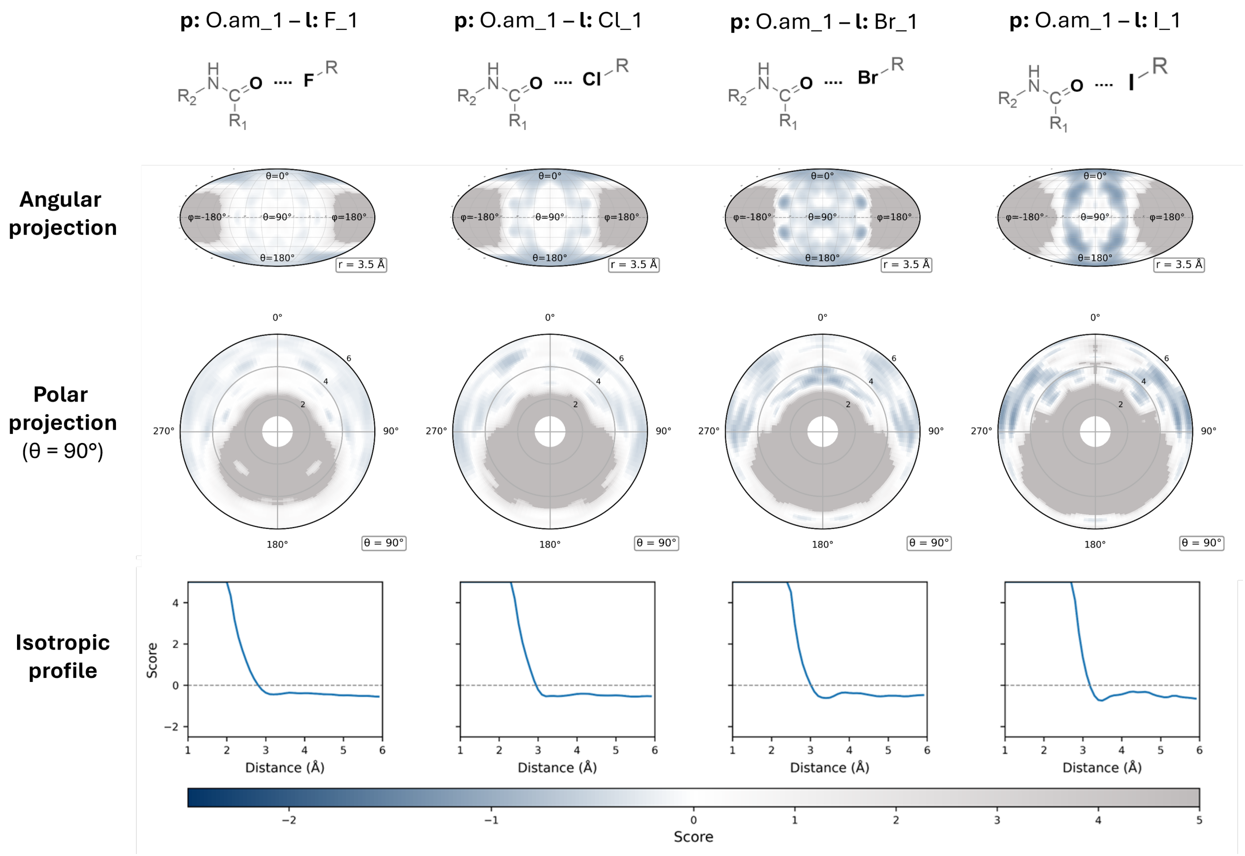


Figure S5: **Anisotropic knowledge-based potential scores for four halogen-bond interaction types with an amide oxygen.** Each column corresponds to a distinct interaction pair, identified by internal atom typing names and schematic depictions (top row): amide O interacting with F, amide O interacting with Cl, amide O interacting with Br, and amide O interacting with I. The second row shows Mollweide projections of the score mapped onto the spherical surface at a fixed radial distance from the protein atom (r = 3.5 Å for all columns), revealing the directional preferences of each interaction. The third row displays polar plots of the score within the azimuthal plane ($\theta = 90$ °), with the radial axis corresponding to the protein–ligand distance (Å). The bottom row shows the isotropic radial score profile from DESPOT-iso. Negative scores (blue) indicate geometries observed more frequently than expected from a reference distribution, while positive scores (gray) indicate disfavored geometries. Progressing from fluorine to iodine, the interaction profiles show increasingly pronounced angular localization perpendicular to the amide plane, consistent with the growing strength and directionality of halogen bonding with increasing halogen polarizability.
